# Priority effects in herbivore communities vary in effect on plant development and reproduction in four Brassicaceae plant species

**DOI:** 10.1101/2022.12.10.519923

**Authors:** Daan Mertens, Jacob C. Douma, Bram B. J. Kamps, Yunsheng Zhu, Sophie A. Zwartsenberg, Erik H. Poelman

## Abstract

1. Insect herbivores can directly affect plant reproduction by feeding on reproductive tissues, or indirectly by feeding on vegetative tissues for which plants are unable to compensate. Additionally, early arriving herbivores may have cascading effects on plant fitness by altering the richness and abundance of the later arriving community.
2. Studies are divided on whether herbivory early in the development of plants can impact plant fitness and whether these effects are predominantly mediated through changes in plant development or subsequent herbivory. Obtaining insight into the generality and consistency of mediated effects on plant reproduction induced by early-season herbivory requires a comparative approach across plant species and environmental conditions.
3. By excluding the herbivore community in an exclosure experiment and by manipulating early-season herbivory in a common garden experiment replicated across four Brassicaceae species and two years, we tested whether early-season herbivory could affect plant development, reproduction, and the herbivore communities associated with individual plants. In addition, we tested a causal hypothesis to disentangle the direct effect of herbivory on plant reproduction, and their indirect effect through changes in the development of plants.
4. Early-season herbivory affected plant development and reproduction, but effects were highly dependent on the plant species, the inducing herbivore species, and the biotic and abiotic environment. The exploratory path analysis indicated that plant reproduction was best predicted by variation in plant development, explaining up to 90.88% of the total effect on seed production. Even though the richness and abundance of the subsequent herbivore community were conditionally affected by the initial colonising herbivore, herbivore pressure is predicted to have only minor effects on reproduction. Importantly, the effects of herbivore pressure on seed set were not direct but were mediated by changes in plant development.
5. *Synthesis:* Early-season herbivory has the potential to affect plant reproduction through changes in the development of plants and, less strongly, through effects mediated by the plant-associated herbivore community. However, as plants are often able to compensate and attenuate the effects induced by herbivory, the detection, sign, and strength of effects are highly dependent on the plant species and environment.

## Introduction

Over their lifetimes, plants interact with a diverse and highly dynamic community of insect herbivores. As the costs of herbivory are dependent on the plant’s ontogeny and environment, plants may not always be able to fully compensate for the consumptive effects of herbivores, resulting in reduced reproductive output (Boege & Marquis 2005; Lehndal & Agren 2015; Ochoa-Lopez *et al*. 2015). Herbivores directly affect plant reproduction by feeding on reproductive tissues or predating on seeds, or indirectly by feeding on vegetative tissues important for the growth and development of plants. These indirect effects on reproduction mediated by changes in the plant’s development may be amplified by other ecological processes, such as competitive interactions with neighbouring plants (Kolb *et al*. 2007; de Vries *et al*. 2017).

However, the indirect effects of herbivory on plant reproduction need not be exclusively mediated by the plant itself. Herbivore-induced changes in plant phenotype have been shown to affect the plant’s susceptibility to later attacks, connecting plant herbivores through a network of indirect interactions (Ohgushi 2005; Kessler & Halitschke 2007; Huang *et al*. 2017). These changes in the phenotype of plants emerge through induced plant responses involvying myriad chemical and morphological traits to enhance resistance to the attacker (Karban 2011; Schuman & Baldwin 2016), or result from antagonists manipulating plant responses or modifying the plant tissues on which they feed for their own benefit (Lill & Marquis 2003; Behmer 2009; Dussourd 2017). In addition to alterations in the identity of attackers, induced plant responses that occur early in the season can affect the trajectory of insect population growth that persists throughout the development of the plant (Karban 1993; Wold & Marquis 1997). Although direct interactions with herbivores are often highly transient, priority effects caused by individual attackers may thus influence the subsequent interactions plants are exposed to, even after the causal biotic interaction ceases to persist (Wurst *et al*. 2015; Han *et al*. 2020).

These priority effects in which early colonisers affect the trajectory of community assembly are increasingly recognised as potential drivers of ecological and evolutionary dynamics (Poelman & Kessler 2016; Kafle *et al*. 2018; Mertens *et al*. 2021a). A growing number of studies identified priority effects of transient herbivory on either above or belowground herbivore communities (Barber *et al*. 2012; Stam *et al*. 2018). These effects can cascade through the carnivore community (Hernandez-Cumplido *et al*. 2016), the fungal community (Kostenko *et al*. 2012; Abdala-Roberts *et al*. 2019), the pollinator and florivore community (Hoffmeister *et al*. 2016; Chauta *et al*. 2017; Rusman *et al*. 2020), and can ultimately affect plant reproduction (McArt *et al*. 2013; Machado *et al*. 2018; Stam *et al*. 2019; Rusman *et al*. 2020). The ecological costs plants suffer through altered interactions with antagonists and mutualists can accumulate throughout the plant’s lifetime, with indirect effects possibly outweighing the direct (i.e. consumptive) costs of the initial attack on plant fitness (Poelman & Kessler 2016).

The notion that transient herbivory can impact plant fitness through both community-mediated effects and changes in plant development reveals a potential key role for early-season herbivores in driving ecological and evolutionary dynamics. Their effects on reproduction can be modulated by their priority effects on the assembly of the subsequent community, as well as through changes in the plant’s development ensuing from the damage to valuable plant tissues early in the development of plants. In addition, plant responses to initial herbivory and the induced changes in phenotype and development of the plant are determined by functional traits such as the feeding guild and host specialisation of the early-season herbivore and are likely to be important in determining the emergent effects on subsequent plant development, interactions with the community, and plant reproduction (Ali & Agrawal 2012; Mertens *et al*. 2021b; de Bobadilla *et al*. 2022).

Disentangling the direct and indirect effects of early-season herbivores on plant reproduction is challenging, as direct effects of herbivores on plant performance often include an induced response that results in altered interactions with the plant-associated community and indirect effects on plant performance (Gruntman & Novoplansky 2011; West & Louda 2018). Moreover, while indirect effects induced by herbivores are increasingly documented across different study systems, it remains unknown whether similar conditions in the environment cause specific herbivore species to induce similar effects in related plant species. A comparative approach across plant species would provide insight into the generality and consistency of plant- and community-mediated effects on plant reproduction and increase our understanding of the trade-offs in life history strategies plants face.

In this study, we tested the effects of early-season herbivory on plant reproduction in four closely related annual Brassicaceae species replicated in two growing seasons (2017 and 2018), and evaluated whether these effects were comparable between plants grown in an open-field common-garden and plants grown in tents (i.e. excluding effects mediated by the herbivore community). To test the effect of early-season herbivory, we manipulated the presence of either *Myzus persicae* aphids or *Pieris rapae* caterpillars on plant seedlings, or left plants untreated. We investigated **i)** whether early-season herbivory affected plant female reproduction and whether effects were dependent on the presence of the herbivore community, **ii)** whether we could formulate and validate a causal hypothesis in the form of a path analysis describing community-mediated and plant-mediated effects on plant reproduction, **iii)** whether early-season herbivory affected the variables used in this path model, and **iv)** whether early-season herbivory affected the composition or structure of the subsequent herbivore community associated with individual plants. Our results provide novel insights into the relative importance of direct and indirect interactions in plant-insect communities, and we discuss its implications for the evolution of plant defence strategies.

## Materials and methods

### Study system

We tested the effects of early-season herbivory on plant reproduction, plant development, and the herbivore communities associated with four annual Brassicaceae species: *Brassica nigra* W.D.J. Koch, *Raphanus raphanistrum* L., *Sinapis arvensis* L., and *Rapistrum rugosum* (L.) All. (Table S1). These species are annual herbaceous plants native to the Netherlands, have similar ecological niches, are overlapping in their development and phenotype (e.g. *S. arvensis* is shorter lived than the other plant species and *B. nigra* typically grows taller than the other plant species), and share a substantial part of the community of herbivore species (Mertens *et al*. 2021c). For each plant species, we collected seeds from at least 25 mother plants propagated by open pollination at the experimental fields of Wageningen University, The Netherlands (51°59’26.5”N; 5°39’50.5”E). Seeds were sown in trays with potting soil (Lentse Potgrond) and germinated in a glasshouse. After germination, plants were transplanted to peat soil cubes. Three-week-old plants were placed under a roofed shelter to acclimatise them to field conditions. Plants were transplanted to the experimental field when they were four weeks old (mid-May; week 21 of 2017 and 2018). Adult wingless *M. persicae* (green peach aphids) and neonate *P. rapae* caterpillars (small cabbage white) used in our experiment originated from stock cultures kept under greenhouse conditions (22 ± 1°C, 50-70% r.h., L16:D8) at the Laboratory of Entomology, Wageningen University. The aphids were reared on *Raphanus sativus* (radish) and caterpillars were reared on *B. oleracea* var. *gemmifera* cv. Cyrus (Brussels Sprouts).

### Common garden set-up

Two days after planting in the open field, we infested plants with either three adult wingless aphids, two first-instar caterpillars, or left plants untreated. The 12 possible combinations of four plant species and three treatments were randomly assigned to plots consisting of nine plants in monoculture planted one meter apart. Each of the 12 combinations was replicated eight times, resulting in 96 plots in each field season. Plots measured 3m x 3m and were separated from each other and the field edge by 4-meter-wide grass lanes. To obtain edge uniformity, we planted a strip of *B. nigra* in high density (six plants per square meter, 1m wide) around the experimental field. In addition, we installed a mesh fence and kites to prevent herbivory by vertebrates.

We recorded the development of herbivore communities on the five central plants in each plot (excluding the four corner plants) by monitoring individual plants from seedling to seed set. In cases where a plant died before the second monitoring round, we monitored one of the corner plants instead. Recording of the herbivore communities started two days after the application of the treatments. We monitored the development of herbivore communities on individual plants by weekly counts early in the season and by biweekly counts later in the season. Insects were identified *in situ* to species or family level. If accurate identification was not possible, we included the observations as morphospecies in our data (Table S2). In addition to observations of the herbivore community, we recorded a set of plant parameters as proxies for plant biomass and development: plant height (measured from the ground to the top of the plant), diameter (measured as the distance between the two most distal leaves), the number of true leaves, and the number of flowering and seed-carrying branches (aggregated as reproductive branches). The height and diameter of plants were used to derive the volume of a cone, representing plant biomass as a single volume parameter in the subsequent analyses. Finally, to assess the effects of early-season herbivory on plant reproduction, we measured the seed set of the plants as a proxy for plant fitness. We harvested all plants monitored in the experiment after the start of plant senescence but before the siliques started losing seeds. For each plant, the total number of seeds was estimated by extrapolating the weight of 100 seeds to the total seed biomass and rounding to the nearest natural number.

### Herbivore exclosure set-up

To assess the effects of early-season herbivory on plant reproduction while excluding effects mediated by the herbivore community, we installed an exclosure experiment that ran in parallel with the open field experiment. Plants were planted in mesh cages (measuring 4m length x 4m width x 2.5m height), with nine plants in each tent planted at 1m equidistance. Mesh size of the tents was 0.6mm, which excluded all arthropods while allowing light and air into the tent. Mesh cages were located at the same experimental site as the common garden experiment. Treatments were applied by placing either three adult wingless aphids or two first instar caterpillars on a fully developed leaf and enclosing this leaf in a mesh bag. To control for a possible effect of enclosing leaves, we also enclosed one of the leaves of plants that were left untreated. After 14 days we removed the treatment by excising the bagged leaf with a razor. Bagging the herbivore infested leaf enhanced the chance of keeping the tent free of herbivores after the induction treatment. To ensure pollination, we placed commercially available *Bombus terrestris* hives (*Natupol Smart*; Koppert Biological Systems; Rodenrijs, The Netherlands) and *Lucilia sericata* fly pupae (*Natupol Fly*; Koppert Biological Systems; Rodenrijs, The Netherlands) in the cages at the start of flowering. The number of replicates in the exclosure experiment was year-dependent due to unintentional infestations of plants and the limited number of mesh cages available (Table S1). Plant reproduction was estimated using the same protocol described for the common garden experiment.

## Statistical analysis

As we directly manipulated the presence of *P. rapae* and *M. persicae* as part of our experiment, we excluded these herbivore species from the herbivore community observations. In addition, we removed plants for which the infestation treatment was considered unsuccessful (the insects applied as treatment were not observed in any of the two first observation rounds) from our dataset and removed plants that were monitored less than four times during the field season (Table S1). Analyses were carried out in R (R Core Team 2014) using the *lme4* (Bates *et al*. 2015), *nlme* (Pinheiro *et al*. 2012), *PiecewiseSEM* (Lefcheck & Freckleton 2015), *ggplot2* (Wickham 2009), *emmeans* (Lenth *et al*. 2019), *mgcv* (Wood 2011), *gamm4* (Wood *et al*. 2017), *Multcomp* (Hothorn *et al*. 2008), *car* (Fox & Weisberg 2018), *vegan* (Oksanen *et al*. 2012), *BiodiversityR* (Kindt & Coe 2005), and *bipartite* (Dormann *et al*. 2008) packages.

### Effect of early-season herbivory on plant reproduction

To test whether early-season herbivory affected plant reproduction, we analysed the square-root-transformed number of seeds for each of the plant species separately using mixed effect models. Models included treatment, year of the field season, and their interaction as explanatory variables and included plot identity or identity of the mesh cage as random intercept. As seed set of plants in the open field was not directly comparable to that of seeds produced by plants in the tents, we analysed the common garden and herbivore exclosure experiments separately. In each analysis, we formulated a set of models to include either a variance structure that was year-dependent, treatment-dependent, or dependent on both factors combined, or assumed homogeneity of variance among all factor levels (Bates *et al*. 2015). We selected the model with the lowest Akaike Information Criteria (AIC) score and used diagnostic plots to verify that model assumptions were met (Johnson & Omland 2004; Zuur *et al*. 2009; Harrison *et al*. 2018). We estimated the effect size and significance of fixed factors using type II Wald χ2-tests. Pairwise post-hoc comparisons among treatments were performed for each year separately using Tukey’s honest significant difference test.

### Path analysis

To disentangle the direct and indirect effect of herbivores on plant fitness, we formulated causal models that relate plant performance and herbivory early in the season to plant performance and herbivory later in the season, and ultimately plant reproduction at the end of the growing season. Path analysis is a multivariate technique that allows modelling the multivariate dependency between variables based on a priori causal hypothesis (Shipley 2016). Once such a hypothesis is consistent with the data, it can be used to quantify the direct and indirect effects of herbivory and plant development on reproduction. We formulated a set of path models describing our initial causal hypothesis (Fig. 1). These path models differed in how they included year and plant-species effects on the intercept of variables in the path model while constraining the path coefficients to be the same across year and plant species, i.e. the strength of the relationships between the variables (Table S3). To avoid an overparameterization of the path model, we aggregated parameters that were repeatedly measured over the growing season, i.e. the volume of plants, the number of leaves, and the abundance and richness of the folivore community, into early-season (observation rounds 1 and 2; weeks 1 and 2 after the start of the experiment in both 2017 and 2018), mid-season (rounds 3 and 4; between week 3 and 5 in 2017 and between week 3 and 4 in 2018), and late-season (rounds 5 and 6; between week 6 and 8 in 2017, and between week 5 and 7 in 2018) proxies by taking the maximum observed parameter value within each subdivision. All variables in the path analysis were checked for normality and transformed when normality was violated. The volume of plants, the number of reproductive branches, and the number of seeds were square-root-transformed, and the abundance of herbivores associated with leaves, and the abundance of florivores and seed predators were transformed using a log(x+1) transformation. Variables were then centred and standardized to a mean value of 0 and a standard deviation of 1. The path model was constructed by specifying linear mixed models (LMM) including the identity of plots as random effect, and accounting for the heterogeneity of residuals by using variance functions when resulting in a better model fit based on AIC. The path model was then evaluated by AIC and Fisher’s C global goodness of fit statistic with the associated P-value, where P > 0.05 indicates that the data are sufficiently well represented by the path model (Shipley 2009; Lefcheck & Freckleton 2015). To test the validity of the causal assumption when relaxing the constrain on path coefficients, we fitted the causal model to each plant species, the two years, and all combinations of plant species and year separately (Table S3). Model parsimony when either retaining or relaxing the constraint on path coefficients was evaluated using AIC (Shipley & Douma 2020; Douma & Shipley 2021).

**Figure 1:**
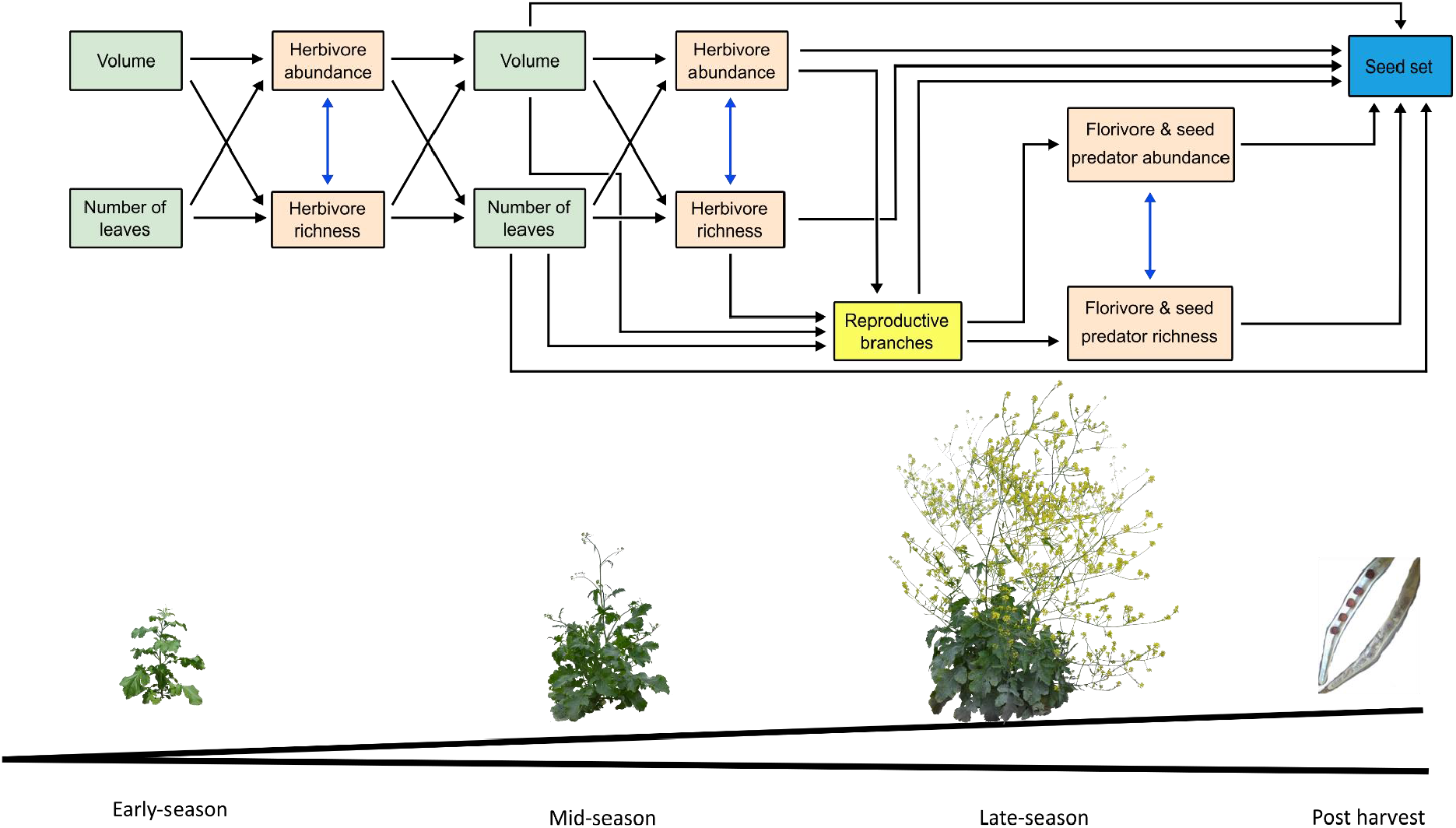
Path diagram of the causal hypothesis expressing the relation between the measured proxies for plant biomass and the observed herbivore community early in the plant development, plant biomass and the observed herbivore community in the middle of the plant growing season, the number of reproductive branches and associated florivores and seed predators late in the season, and plant reproduction. Black arrows represent unidirectional relationships among variables while double-headed blue arrows show correlated errors. Variables related to plant development, the observed herbivore community, the number of reproductive branches, and seed production are represented in green, red, yellow, and blue rectangles, respectively.

As our initial causal hypothesis was not supported by the data (Table S3), we proceeded to an exploratory approach by reformulating path models that were found to be consistent with the data. We removed all paths in our initial hypothesis which were not supported and included all paths for which we found a significant independence claim and, importantly, whose causality could be supported by ecological knowledge. Variables that showed dependence but between which we could not presume a biologically sound causal relation were incorporated as correlated error. The final structure and fit of the optimized model depended on how the effects of plant species and year on the estimations of variables in the model were included in the path model (Table S4). The most parsimonious path model was selected based on AIC (Shipley & Douma 2020).

To determine the relative importance of variables related to plant development and those related to the herbivore community in their effect on plant reproduction, we evaluated the direct and indirect effect of each variable in the model on seed set. Direct effects are represented by the direct path from the variable of interest to seed set. An indirect effect represents the effect of the variable of interest to seed set mediated by another variable, and can be calculated as the product of the path coefficients along the path. The total indirect effect is the sum of all possible indirect effects. We used standardized path coefficients to allow comparison of effects among the different variables (Shipley 2016). Finally, we fitted the optimized exploratory path model on the different year-by-species combinations to assess the generality of the model structure and consistency of direct and indirect effects on seed production across different plant species and the two years.

### Effect of early-season herbivory on path variables

To estimate the degree to which early-season herbivory could affect plant fitness, we fitted univariate regressions for all variables in the path model (mixed effect models). All regressions included treatment, year of the field season, and their interaction as explanatory variables and included plot identity as random intercept. For each of the path variables tested, we formulated a set of models differing in how they accounted for heterogeneity of variance among factor levels and selected the model with lowest AIC. We estimated the effect size and significance of fixed factors using type II Wald χ2-tests. Pairwise post-hoc comparisons were performed for each year separately using Tukey’s honest significant difference test.

To make a tractable path model, the herbivore community was aggregated to early-, mid-, and late-season proxies. To test the robustness of this choice, we performed additional analyses on the unaggregated form of variables which were repeatedly measured while controlling for the time since the start of the experiment. A preliminary analysis of the relationship between each of the repeatedly measured variables (herbivore richness and abundance, and plant volume and number of leaves) and the day after the start of the experiment indicated a nonlinear relationship between time and the response variable. We therefore applied Generalized Additive Mixed Models (GAMM) with Gaussian, gamma, or negative binomial probability distribution and thin plate regression splines to assess dynamics over time (Zuur 2012; Wood 2017). These models included the treatment, year of the field season, and their interaction as fixed factors and included plant identity nested within plot identity as random intercepts to account for the dependency of observations and repeated measurements on the same plant. We then compared models fitted with a single smoothing function estimating the relation between the response variable and day after start of the experiment, with models that fitted a different smoothing function for each early-herbivory treatment level (three smoothers), year of the field season (two smoothers), or the interaction between treatment and year (six smoothers) using likelihood-ratio tests. The effect size and significance of variables in the model were estimated using likelihood-ratio tests. As we found strong variation across years relative to the effects of our treatments, we repeated the analysis for each field season separately.

### Effect of early-season herbivory on herbivore community composition and structure

In a final set of analyses, we explored whether the composition and structure of herbivore communities differed among plant species and treatments in each of the two years. To visualise the herbivore community observations, we constructed a bipartite interaction network linking herbivore species and plant species and used multivariate ordinations (non-metric multidimensional scaling with three dimensions) of the herbivore community on individual plants. We assessed community composition (i.e. incidence of herbivore species) based on the Sørensen dissimilarity matrix, and community structure (composition weighted for species abundances) by calculating the Euclidean distance of Hellinger-transformed cumulative abundance data for each plant individual (Legendre & Gallagher 2001). Using these distances, we tested for differences among differently treated plants of the same plant species through a non-parametric permutational multivariate analysis of variance with 1000 permutations (PERMANOVA) (Anderson 2001). To ensure valid permutation of communities, we specified the dependency of observations on plants in the same plot in a stratified permutational design. As this permutation design requires an equal number of samples in each permutable group, we randomly sampled three plants per plot to obtain an equal sample size and then performed the stratified PERMANOVA analysis. This procedure of randomisation followed by PERMANOVA was repeated 1000 times and we report the median, first and third quantiles, and 5th and 95th percentiles of the results obtained by repeated permutation analysis.

## Results

### Effect of early-season herbivory on plant reproduction

Effects of early-season herbivory on plant reproduction strongly depended on whether plants were grown in the exclosure set-up or in the open field. In the community exclosure experiment, we found that early-season herbivory affected plant reproduction (Table 1), although the sign (i.e. positive or negative) and strength of the effect was strongly dependent on the plant species and year. Interestingly, inducing herbivores could have a positive effect on seed production compared to non-treated plants in one plant species and a negative effect on seed production in another plant species or year (Fig. 2). For example, in the 2017 growing season, we found that early-season herbivory by aphids on *R. raphanistrum* resulted in a 54% decrease in the square-root-transformed number of seeds produced relative to untreated plants, while in *S. arvensis* it resulted in a 64% increase in seed production. Similarly, the sign of the effect induced by a specific early-season herbivore was year-dependent. For example, in 2017 we observed that early season herbivory by caterpillars feeding on *B. nigra* resulted in a 43% reduction in seed set, while in 2018 herbivory by caterpillars resulted in a 91% increase in seed production compared to plants which were left untreated. While the effects of the inducing herbivore in the exclosure experiment could be substantial, we did not find any evidence for an overall effect of early-season herbivory on seed production for plants grown in the common garden set-up (Table 1). This was confirmed by the post-hoc analysis, showing that only the pairwise difference between untreated *R. rugosum* plants and plants challenged by caterpillars in the 2018 season was significant (1 out of 48 pairwise comparisons). Overall, these results indicate that the effect of early-season herbivory on plant reproduction was strongly dependent on the inducing herbivore species, the mediating plant species, the biotic environment (i.e. exclusion of the community or open field) and the abiotic environment (illustrated by the year effects in the exclusion experiment) in which plants grew.

**Table 1.** Overview of effect size of early-season herbivory (Treatment), year of the field season (Year), and their interaction (Treatment*Year) on square-root transformed number of seeds produced by the different plant species in the community exclosure and open field experiments. Models were adjusted to account for heterogeneity of model residuals using a variance function across different treatments, years, all factor levels, or were not adjusted when no heterogeneity found (indicated as none). Values in bold indicate significant factor effects (p < 0.05).

| Experiment | Plant species | Variance function | Explanatory variable | $\chi^2$ | df | p-value |
| --- | --- | --- | --- | --- | --- | --- |
| Enclosure | *Brassica nigra* | none | Treatment | 3.6347 | 2 | 0.1625 |
|  |  |  | Year | 31.5466 | 1 | **< 0.0001** |
|  |  |  | Treatment*Year | 19.613 | 2 | **< 0.0001** |
|  | *Raphanus raphanistrum* | Treatment | Treatment | 25.609 | 2 | **< 0.0001** |
|  |  |  | Year | 0.0574 | 1 | 0.8107 |
|  |  |  | Treatment*Year | 6.8773 | 2 | **0.0321** |
|  | *Sinapis arvensis* | Treatment*Year | Treatment | 6.6843 | 2 | **0.0354** |
|  |  |  | Year | 3.1211 | 1 | 0.0773 |
|  |  |  | Treatment*Year | 3.2576 | 2 | 0.1962 |
|  | *Rapistrum rugosum* | none | Treatment | 5.8281 | 2 | 0.0543 |
|  |  |  | Year | 0.4815 | 1 | 0.4877 |
|  |  |  | Treatment*Year | 1.2685 | 2 | 0.5303 |
| Open field | *Brassica nigra* | Treatment*Year | Treatment | 1.2456 | 2 | 0.5364 |
|  |  |  | Year | 2.1927 | 1 | 0.1387 |
|  |  |  | Treatment*Year | 2.5783 | 2 | 0.2755 |
|  | *Raphanus raphanistrum* | Treatment*Year | Treatment | 0.4965 | 2 | 0.7802 |
|  |  |  | Year | 0.1992 | 1 | 0.6554 |
|  |  |  | Treatment*Year | 0.0223 | 2 | 0.9889 |
|  | *Sinapis arvensis* | Year | Treatment | 1.7279 | 2 | 0.4215 |
|  |  |  | Year | 65.7476 | 1 | **< 0.0001** |
|  |  |  | Treatment*Year | 1.6954 | 2 | 0.4284 |
|  | *Rapistrum rugosum* | Treatment*Year | Treatment | 3.7083 | 2 | 0.1566 |
|  |  |  | Year | 0.2349 | 1 | 0.6279 |
|  |  |  | Treatment*Year | 3.5473 | 2 | 0.1697 |

**Figure 2.**
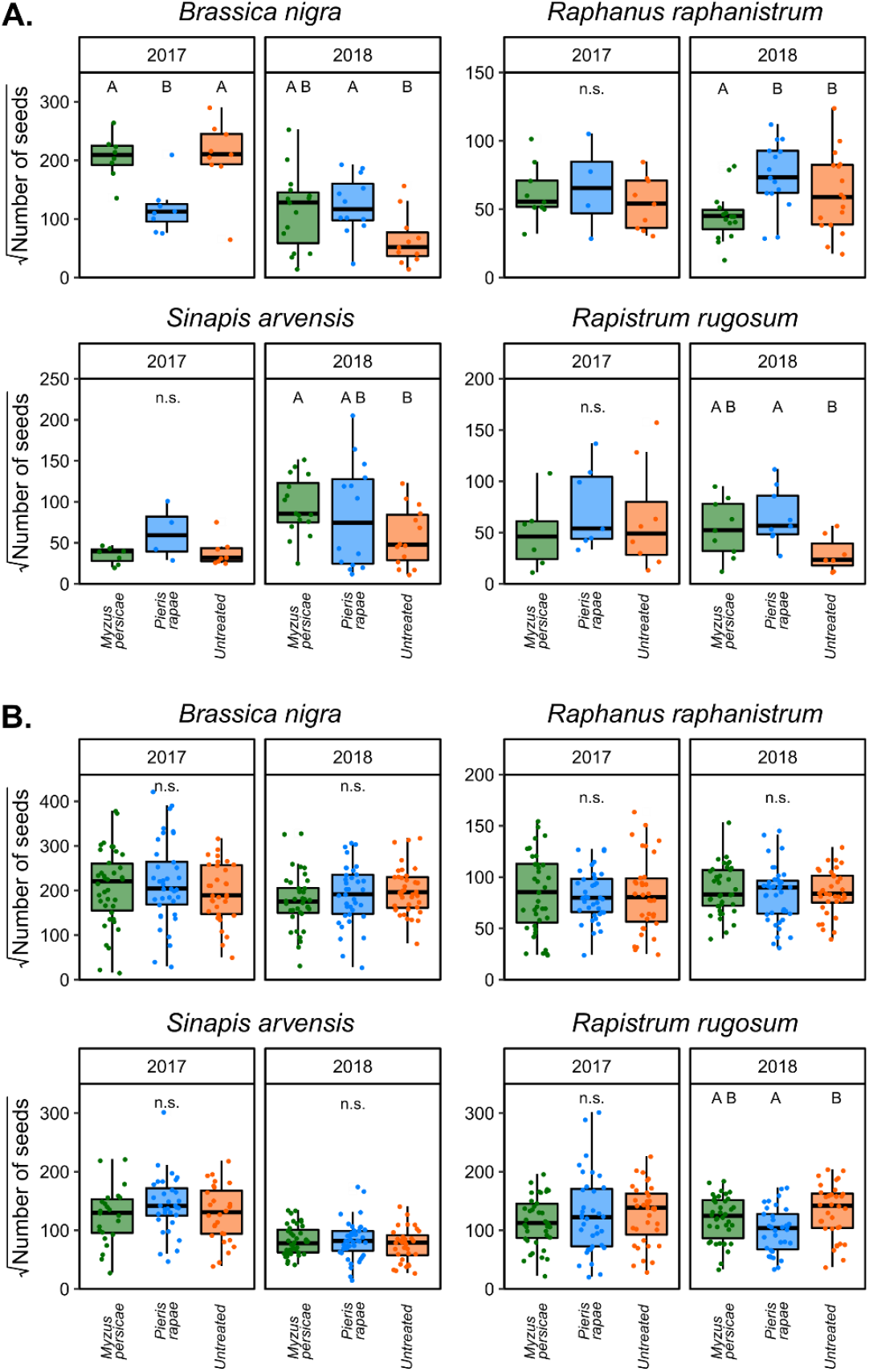
Seed production by plants belonging to four different plant species measured in two field seasons (2017 and 2018), **A**. when the herbivore community was excluded, and **B**. when plants could interact with their full associated communities. Plants or were challenged early in the season by *Myzus persicae* aphids (green) or *Pieris rapae* caterpillars (blue), or were left untreated (orange). Pairwise comparisons were performed separately for each plant species by field season combination. Boxplots show median, first and third quantiles, and 95% interval. Dots represent individual observations. Boxes within each panel with no letters in common are significantly different based on Tukey’s HSD (p < 0.05), whereas n.s. indicates no significant differences were found.

### Path analysis

Exploration of path models fitted on the full data or subsets of the data based on combinations of the plant species and field seasons, revealed that our *a priori* hypothesis expressing the relation between the measured proxies for plant biomass, the observed folivore herbivore community, the number of reproductive branches and associated florivores and seed predators, and seed production, was not supported by the data (Table S3). When using an exploratory approach to formulate a causal hypothesis, several changes were made compared to our initial model (cf. Figs 1 and 3). First, causal relationships between early-season biomass and mid- or late-season variables to seed production had to be added, suggesting alternative indirect paths through which early-season variables can affect plant reproduction (Fig. 3). For example, early season biomass in terms of plant volume, as well as the number of leaves had a direct and significant causal relation with the number of reproductive branches plants produced. Plants with more leaves early in the season produced more reproductive branches, while plants with a higher volume early in the season produced fewer branches when controlling for indirect effects. Second, the exploratory analysis suggests that early-season herbivore abundance, but not richness, had a positive effect on plant reproduction, while mid-season herbivore richness, but not abundance, had a negative effect on reproduction (Table 2). Importantly, these effects were exclusively mediated by variables related to plant development, with neither foliar herbivory nor herbivory of reproductive tissues having a direct effect on seed production. In addition, the causal effects of herbivory on variables related to plant development were estimated to be small. Third, the volume of plants early in the season directly (negatively) affected seed set. The direct effect of early-season variables not mediated through volume and leaf number later in the season suggests that their effects may be mediated by variables not included in the analysis (e.g. variables related to plant ontogeny). Taken together, the exploratory model fitted on all data independent of plant species and year of the field season indicated that plant development had the strongest causal relations with seed production, while effects induced by herbivory were less strong (Table 2). Plant characteristics accounted for 90.88% of the total effects on seed production (74.22% direct, 16.66% indirect) while herbivore pressure accounted for 9.12% of the effects on seed production (0.00% direct, 9.12% indirect).

**Table 2.** Overview of the direct, indirect, and total effect of variables included in the optimized exploratory piecewise path model on seed production of plants in the open field experiment. Variables are further annotated as related to plant development and biomass (P) or related to herbivore pressure (H). The volume of plants and the number of reproductive branches was square-root-transformed, and the abundance of herbivores, florivores, and seed predators was transformed using a log(x+1) transformation. Effects larger than 0.1 are indicated in bold.

| Time | Variable | Type | Direct | Indirect | Total |
| --- | --- | --- | --- | --- | --- |
| Early season | Number of leaves | P | 0.00 | 0.07 | 0.07 |
|  | Volume | P | -0.08 | 0.03 | -0.05 |
|  | Herbivore richness | H | 0.00 | 0.00 | 0.00 |
|  | Herbivore abundance | H | 0.00 | 0.07 | 0.07 |
| Mid-season | Number of leaves | P | **0.19** | 0.00 | **0.19** |
|  | Volume | P | **0.39** | 0.02 | **0.42** |
|  | Herbivore richness | H | 0.00 | 0.00 | 0.00 |
|  | Herbivore abundance | H | 0.00 | 0.00 | 0.00 |
| Late season | Reproductive branches | P | 0.05 | 0.00 | 0.05 |
|  | Florivore & seed predator abundance | H | 0.00 | 0.00 | 0.00 |
|  | Florivore & seed predator richness | H | 0.00 | 0.00 | 0.00 |

**Figure 3.**
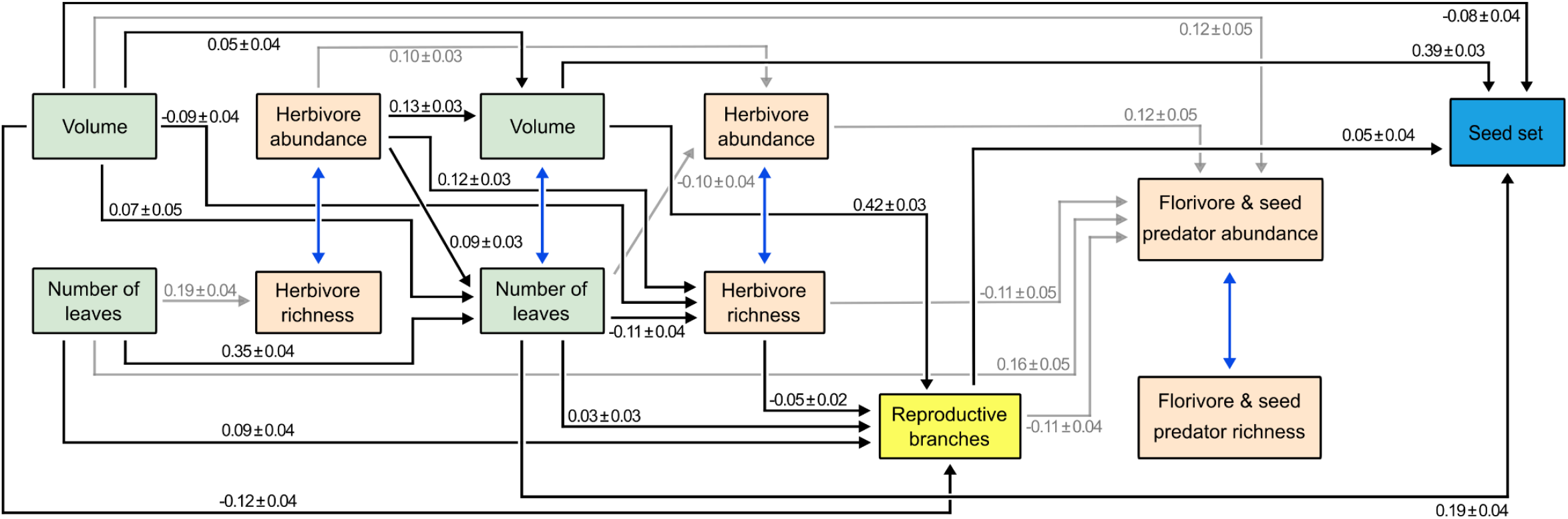
Exploratory piecewise path model obtained by optimizing our causal hypothesis where path coefficients and estimation of variables were constrained to be equal across all plant species and the two years. Optimization was performed by removing all paths in our initial hypothesis which were not supported by the data and including all paths for which we found a significant independence claim. Relations between variables for which we could not presume a biologically sound causal relation were included by their covariance. Black arrows represent direct and indirect paths between variables and seed production, grey arrows show paths that were included in the path model but do not affect reproduction, and blue double-headed arrows indicate relations between variables that were included as correlated error. Arrows are annotated by their standardised path estimate and standard error. Variables related to plant development, the observed herbivore community, the number of reproductive branches, and seed production are presented in green, red, yellow, and blue rectangles, respectively.

Fitting the optimised exploratory piecewise path model to different subsets of the data to remove equivalence constrains on path coefficients revealed that the estimation of path coefficients strongly depended on the plant species, the year, or the plant species by year combination (Table S5). The estimated path coefficients obtained when fitting the model on subsets of the data frequently fell outside of the 95% confidence interval around the estimation of the same coefficient obtained when fitting the path model on the full data (Table S6). The dependence of path coefficients on the year and plant species had substantial influence on the estimation of direct and indirect effects on seed set (Table S7). However, plant development remained more important in predicting plant reproduction than variation in the richness or abundance of the herbivore community. Even though the estimated values of path coefficients depended on the subset of the data to which the model was fitted, constraining path coefficients to be equal among plant species and the two years was found to result in the most parsimonious model based on AIC (path coefficients fully constrained: AIC = 157; equivalence constraint removed between years but not plant species: AIC = 325; equivalence constraint removed between plant species but not between years: AIC = 685; equivalence constraints on path coefficients removed for both year and plant species: AIC = 1329). However, this generalisation may be the result of the increased uncertainty in the path coefficients with the smaller sample sizes of each subset. In addition, it is likely that not all path coefficients are dependent on the plant species or year.

### Effect of early-season herbivory on path variables

For most plant species, we found no evidence of significant effects of early-season herbivory (i.e. the induction treatment) on the variables used in the path analysis (Table S8). The exceptions were found for *S. arvensis*, for which our treatments had a significant effect on the mid-season volume of plants (Treatment: χ^2^ = 7.0779, df = 2, p = 0.0290) and had a year-dependent effect on the number of reproductive branches (Treatment * Year: χ^2^ = 7.0812, df = 2, p = 0.0290), and *R. raphanistrum*, where early-season herbivory significantly affected the richness of the herbivore community early in the season (Treatment: χ^2^ = 9.4785, df = 2, p = 0.0087). The overall effects found for *S. arvensis*, but not those found for *R. raphanistrum*, were substantiated by the post-hoc analysis. In 2018, *S. arvensis* plants challenged by aphids had a significantly higher mid-season volume than plants challenged by caterpillars, and in 2017 *S. arvensis* plants treated with aphids produced significantly fewer reproductive branches than caterpillar-treated plants. Even though we did not observe any significant overall effects of our treatments on the variables measured for *R. rugosum*, we did find post-hoc differences between treatments for the mid-season volume of plants and the abundance of florivores and seed predators. *Rapistrum rugosum* plants which were challenged by aphids obtained a significantly lower mid-season volume compared to plants challenged by caterpillars and encountered a significantly higher abundance of florivores and seed predators compared to plants which were left untreated.

We then proceeded with the analysis of the repeatedly measured variables as observed over time, i.e. not aggregated in early-, mid-, and late-season proxies. The models describing a year-specific relation between the different response variables and the day since the start of the experiment generally resulted in the best fit to our data. The only exception was the number of leaves produced by *B. nigra*, for which the change over time was best described by including a different smoother for each treatment (Table S9). When analysing both years simultaneously, our treatments had a significant year-dependent overall effect on the abundance of herbivores on *S. arvensis* (Treatment * Year: χ^2^ = 9.7089, df = 2, p = 0.0078). In addition, early-season herbivory had a significant year-dependent effect on the volume of *R. rugosum* plants (Treatment * Year: χ^2^ = 7.8847, df = 2, p = 0.0194). When analysing the two years separately, the effect of the treatments on the abundance of herbivores on *S. arvensis* was confirmed for the 2018 season (Treatment: χ^2^ = 15.2120, df = 2, p = 0.0005), indicating that plants challenged by aphids early in the season encountered a higher average abundance of herbivores than plants challenged by caterpillars (Fig. S1). We could not confirm that early-season herbivory had a significant overall effect on the volume of *R. rugosum* plants in either year (Fig. S2). In addition, early-season herbivory had a significant effect on the average abundance of herbivores on *R. rugosum* in 2018 (Treatment: χ^2^ = 6.2678, df = 2, p = 0.0436), where plants challenged by aphids early in the season encountered a higher average abundance of herbivores than plants challenged by caterpillars (Fig. S3). Finally, *B. nigra* in the 2017 field season challenged by caterpillars produced a higher number of leaves than plants that were left untreated (Treatment: χ^2^ = 10.1770, df = 2, p = 0.0062) (Fig. S4).

Taken together, these results show that early-season herbivory can affect the richness and abundance of the herbivore community, as well as the plant traits we measured. However, the detectability and strength of effects depends on the plant species and the environmental conditions, as represented by the strong year-dependency of results. Simplifying the repeatedly measured variables by aggregation into early-, mid-, and late-season values obscured the detectability of treatment effects.

### Effect of early-season herbivory on herbivore community composition and structure

Initial exploration of the herbivore communities associated with the four plant species revealed substantial variation in the prevalence of herbivore species over time (Fig. 4). This turnover in the herbivore community in the field corresponded with changes in the ontogeny of plants. The network between the total herbivore community and the different plant species is well connected (i.e. no modularity of interactions between herbivores and specific plant species) and similar across the two years (Fig. 4). Herbivore communities associated with individual plants significantly differed in their composition and abundance-weighed community structure depending on the plant species (results from PERMANOVA composition: Pseudo-F = 37.342, R^2^ = 0.010, df = 3, p = 0.001; structure: Pseudo-F = 61.737, R^2^ = 0.1527, df = 3, p = 0.001) and year (composition: Pseudo-F = 198.230, R^2^ = 0.1769, df = 1, p = 0.001; structure: Pseudo-F = 222.966, R^2^ = 0.1838, df = 1, p = 0.001) (Fig. S5 – S8). Hence, we proceeded to test for treatment effects by analysing each plant species by year separately. The overall effect of our treatments on the composition or structure of herbivore communities was not statistically significant for any of the plant species, and only minimal variation in the composition or structure of the cumulative herbivore community associated with individual plants could be attributed to early-season herbivory (Table 3, Table S10, Fig. S5 – S8).

**Table 3.**
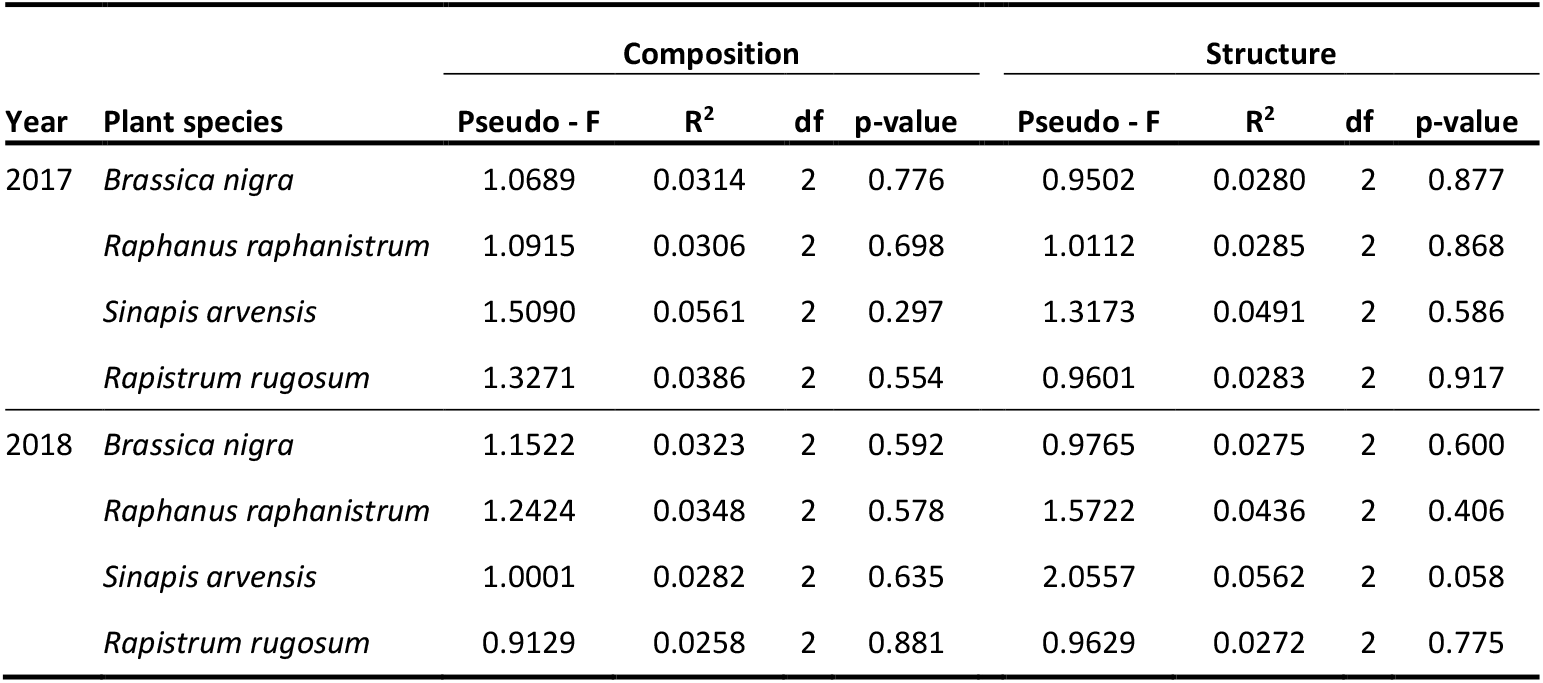
Results of the PERMANOVA analysis testing the effects of early-season herbivory treatments on the composition (incidence) and structure (weighted abundance) of the full herbivore community associated with individual plants in each plant species by year combination. To account for dependency of observations, we applied a stratified permutation design (1000 permutations) with random sampling to ensure equal replication across treatment levels. The table presents the median pseudo-F value with associated R^2^ and statistical significance in terms of p-value. A more complete overview of pseudo-F values per quantile is given in table S9.

| Year | Plant species | Composition |  |  |  | Structure |  |  |  |
| --- | --- | --- | --- | --- | --- | --- | --- | --- | --- |
| | | Pseudo - F | $R^2$ | df | p-value | Pseudo - F | $R^2$ | df | p-value |
| 2017 | *Brassica nigra* | 1.0689 | 0.0314 | 2 | 0.776 | 0.9502 | 0.0280 | 2 | 0.877 |
|  | *Raphanus raphanistrum* | 1.0915 | 0.0306 | 2 | 0.698 | 1.0112 | 0.0285 | 2 | 0.868 |
|  | *Sinapis arvensis* | 1.5090 | 0.0561 | 2 | 0.297 | 1.3173 | 0.0491 | 2 | 0.586 |
|  | *Rapistrum rugosum* | 1.3271 | 0.0386 | 2 | 0.554 | 0.9601 | 0.0283 | 2 | 0.917 |
| 2018 | *Brassica nigra* | 1.1522 | 0.0323 | 2 | 0.592 | 0.9765 | 0.0275 | 2 | 0.600 |
|  | *Raphanus raphanistrum* | 1.2424 | 0.0348 | 2 | 0.578 | 1.5722 | 0.0436 | 2 | 0.406 |
|  | *Sinapis arvensis* | 1.0001 | 0.0282 | 2 | 0.635 | 2.0557 | 0.0562 | 2 | 0.058 |
|  | *Rapistrum rugosum* | 0.9129 | 0.0258 | 2 | 0.881 | 0.9629 | 0.0272 | 2 | 0.775 |

**Figure 4.**
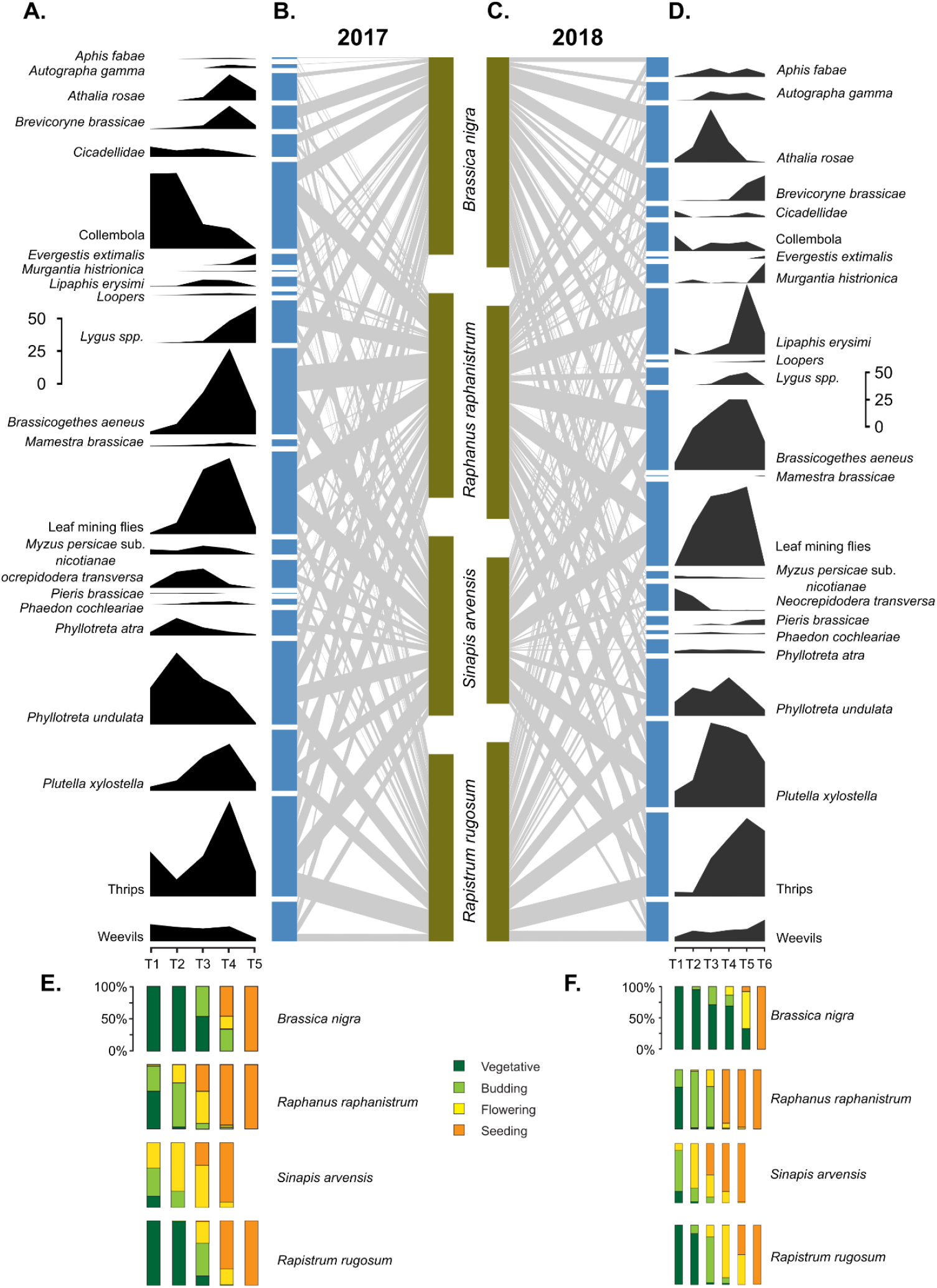
Overview of the number of plants on which each herbivore species occurred (i.e. prevalence) at a given monitoring round (panels A and D) with their respective scales, and the associated interaction network between the herbivore species (blue bars) and plant species (green bars) (panels B and C). Lines connecting herbivore species and plant species in the network represent realised interactions, with the width of these lines representing the frequency of the interaction. The percentage of plants belonging to the different plant species in each stage of ontogeny at a given monitoring round was classified as vegetative (dark green), budding (light green), flowering (yellow), or seeding (orange) (panels E and F). We constructed these figures for 2017 (panels A, B, and E) and 2018 (panels C, D, and F) separately.

## Discussion

We show that early-season herbivory can affect plant reproduction, and that these effects are largely dependent on the plant species, the inducing herbivore species, and the biotic and abiotic environment in which plants grow. When the insect community was excluded, early-season herbivory affected plant reproduction in three out of the four plant species with effects of up to a 6-fold change in seed set (Fig. 2). Effects of early-season herbivory on plant reproduction were attenuated when plants could interact with their full associated community, except for *R. rugosum* plants in 2018, where plants challenged by caterpillars produced significantly less seeds than plants which were left untreated. This was confirmed by an exploratory path analysis of the direct and indirect causal effects between plant development, the herbivore community pressure, and the number of seeds plants produced. Observed levels of herbivore pressure in terms of species richness and abundance of herbivores represented 9.12% of the impact on seed production, and effects were exclusively indirect and mediated by plant development. Variation in plant development and biomass was much more likely to impact seed set (90.88% of the total effect on seed production came from plant biomass and development variables) (Fig. 3, Table 2). Importantly, priority effects induced by early-season herbivory could significantly affect plant development as well as the herbivore community in a year- and plant species-dependent way. However, the effects on variables included in the exploratory model were relatively small, and causal effects induced by variables that were affected by our treatments were often cancelled out by other causal paths involving variables that were unaffected by our treatment, resulting in minimal net effects on plant reproduction. Finally, even though a path model constraining path coefficients to be equal for all plant species and both years was most parsimonious, the strength of causal effects in the exploratory path model differed substantially when fitted to subsets of the data, further emphasising the context dependency of mediated effects. Taken together, our findings provide evidence that priority effects induced by early-season herbivory have the potential to affect plant reproduction through changes in the development of plants and plant-associated herbivore communities, but that the detection, sign, and strength of effects is highly context dependent on the plant species and environment as illustrated by variation in effects across the two years.

Herbivory in life stages when plants are less tolerant to biotic stress can readily be hypothesised to have important and long-lasting consequences for plant development and ultimately plant reproduction. While studies suggest that early-season herbivory has the most substantial effects on plant development, herbivore community assembly, and seed production compared to herbivory in later stages of plant development, these results are not congruent (Garcia & Ehrlen 2002; Adhikari & Russell 2014; Pearse *et al*. 2018; Rusman *et al*. 2020; Rasmussen & Yang 2022). For example, a study on *B. nigra* showed that herbivory by a diverse set of insects in the early ontogenetic stages of plants resulted in a reduced seed set compared to herbivory in later ontogenetic stages (Rusman *et al*. 2020), while a study involving milkweed and monarch caterpillars found that the reproduction of plants challenged early in their development was comparable to that of plants which were left unchallenged (Rasmussen & Yang 2022). Our exclusion experiment highlights the context dependency of effects of early-season herbivory on seed set: In the absence of the plant-associated community, plant reproduction could be increased or decreased compared to plants that were left unchallenged depending on the environmental conditions (i.e. year effects), the plant species, and the identity of the inducing herbivore (Fig. 3). While we cannot fully exclude that abiotic conditions caused by the tents affected seed set, our findings suggest that tolerance to early-season herbivory is likely to be resource-driven or is, non-exclusively, dependent on plant life history strategies in response to specific biotic stressors and abiotic environmental conditions. Variation in selection pressure induced by early-season herbivory may thus arise because plant species or individuals are not equally prepared for, or adapted to, the same stress, or because plants are selected to face different trade-offs over their development which were prevented by excluding the herbivore community (Wright & McConnaughay 2002; Cipollini *et al*. 2014; Lu *et al*. 2016).

When plants were allowed to interact with their full associated community, the effect of early-season herbivory was strongly attenuated, revealing only marginal effects on plant reproduction. The most profound effect of early-season herbivory on plant fitness in a community context was found for *R. rugosum*, where in the 2018 season, plants challenged by *P. rapae* had reduced seed production compared to plants that were left untreated. The attenuated effects of early-season herbivory on plant reproduction in our open field study may be explained by several non-exclusive hypotheses.

First, the pressure induced by our treatment may have been within the tolerance levels of plants, allowing plants to compensate for the damage to tissues and achieve equal reproductive success to that of untreated plants (Strauss & Agrawal 1999; Garcia & Eubanks 2019). This hypothesis is supported by the conditionality in which our treatments affect plant development. For example, in 2018, both *R. rugosum* as well as *S. arvensis* plants challenged by caterpillars had a significantly lower mid-season volume than conspecific plants challenged by aphids, while no significant differences between treatments were found for *B. nigra* or *R. raphanistrum*. This effect was not observed in the 2017 season. Plants may show substantial and context-dependent inter- and intraspecific variability in their ability to compensate for herbivore damage, with outcomes ranging from reproductive or vegetative overcompensation to severe reduction in biomass or seed production (Garcia & Eubanks 2019). Even when plants are unable to fully compensate for the effects induced by early-season herbivory on their vegetative biomass, causal effects on seed production mediated by other factors at different times in the ontogeny of plants may outweigh or cancel out effects mediated by variation in plant development, resulting in minimal net effects of early-season herbivory on plant reproduction (Hambäck *et al*. 2015).

Second, the effects on plant development and reproduction induced by the full plant-associated community may outweigh any effects induced by the initial herbivore treatment. Moreover, in cases where the community, or a subset of key herbivores in the community, is unresponsive to variation in plant traits induced by initial herbivory, the community of herbivores will be homogenised across treatments resulting in comparable effects on plant reproduction (Agrawal 2005; Poelman & Kessler 2016). Even though the effects of prior herbivory by herbivores on the assembly of communities have been documented across a number of study systems (Barber *et al*. 2012; McArt *et al*. 2013; Hernandez-Cumplido *et al*. 2016; Stam *et al*. 2018), several studies report only marginal effects of early-season herbivory on plant-associated communities and reproduction or found no effects at all (e.g. Li *et al*. 2016; Visakorpi *et al*. 2019). The range of ecological outcomes observed across studies is not surprising given the context dependency of herbivore-induced effects and dependency on the plant and herbivore species involved. In support of this hypothesis, our multivariate analysis indicated that herbivore communities associated with the different treatments were not distinguishable in terms of their composition or structure for any of the plant species in either of the two years. We did find priority effects of early-season herbivory on the richness and abundance of herbivores. For example, in 2018, *S. arvensis* and *R. rugosum* plants challenged by aphids early in the season encountered a higher average abundance of herbivores than plants challenged by caterpillars. However, the exploratory path model suggested that the observed variation in herbivore pressure in terms of species richness and herbivore abundance was unlikely to substantially affect seed set, resulting in minimal effects on plant reproduction.

A third hypothesis is that plants were generally not compromised in their responses to differential early-season herbivory. In this perspective, induced responses to initial herbivory are likely selected to incorporate the most likely trade-offs plants face, allowing them to dampen the effects cascading through the associated community (Viswanathan *et al*. 2005; Orrock *et al*. 2015; Karban 2019; Mertens *et al*. 2021a). Plants may fine-tune responses to initial herbivory and the development of functional traits to best fit the most likely or most fitness-impacting future stressors, achieving reproductive success independent of treatment. While a path model assuming equivalence in path coefficients among all plant species and both years was most parsimonious, the substantial variation in path coefficients when fitting the model to the different plant species suggests that the strength of direct and indirect effects linking plant development and herbivore pressure to plant reproduction may not have been generalizable across related plants. Variation among plant species in terms of the strength of plant- or community-mediated effects on seed production may indicate that herbivore pressure varies in how significant it acts as an agent of natural selection driving the evolution of plant life history strategies. In addition, while early-season herbivory may be conjectured to impose frequency-dependent selection on (induced) plant traits and life-history strategies, it is also likely that, for systems with high variability in insect communities over multiple generations, interannual variation in the biotic and abiotic environment leads to stabilised selection in which effects are neutral in their overall pressure on plant trait evolution (Mertens *et al*. 2021a).

A fourth hypothesis relates to the dynamic assembly of herbivore communities in the field. While early-season herbivory initially affects the development and phenotype of the host plant, subsequently arriving herbivores may prefer undamaged or better-developed plants, effectively homogenising herbivore pressure on seed production across different treatments in our experiment (Edwards & Wratten 1983; Rubin *et al*. 2015). Indeed, the notion that herbivores may prefer taller, more apparent plants has been suggested for brassicaceous plants, with potential effects on plant fitness (Schlinkert *et al*. 2016). In addition to host choice in response to variation in plant apparency, the temporal structuring of herbivore communities is intimately connected with the availability of specific niches preferred by herbivores, which in turn is closely related to plant ontogeny (Tonkin *et al*. 2017; Ekholm *et al*. 2020). For example, the strong correlation between the start of flowering and interactions with the pollen beetle *Brassicogethes aeneus* may cause undamaged plants which develop fast, grow tall, and produce abundant inflorescences to attract high numbers of beetles. Feeding damage by these florivores can lead to inflorescences with damaged or undeveloped seeds, and therefore reduced seed production, effectivelly canceling out the headstart these plants had (Williams 2010; Schlinkert *et al*. 2016; Rusman *et al*. 2020).

A fifth hypothesis relates to the fact that this study limits its focus to the associated herbivore community. However, plants maintain ecological interactions with many different community members such as micro-organisms, carnivores, and pollinators which are not assessed in our experiments. These ecological interactions have been shown to be involved in indirect plant defences or tolerance to herbivory and are important in determining plant reproduction (Kos *et al*. 2011; Howard *et al*. 2020). For example, induced responses to early-season herbivory can differentially affect the carnivore community, causing changes in top-down (predator-mediated) control on herbivore communities (Li *et al*. 2016; Lucas-Barbosa *et al*. 2016). Likewise, even though early-season herbivory has been shown to affect flower characteristics (Rusman *et al*. 2019), a diverse pollinator community may ensure pollination by functional complementarity and thus limit herbivore-induced variation in plant reproduction (Santamaria & Rodriguez-Girones 2007).

Finally, the attenuated effect sizes of early-season herbivory in the common garden experiments can be explained by the natural occurrence of the two herbivore species we manipulated. These species are common and readily colonize the plant species used in our experiment under natural conditions (Mertens *et al*. 2021c). Hence, most plants in our experiment interacted with these species at some point in the early- to mid-season stages of their ontogeny. If the timing of attack by the early-season herbivore is not important, we could assume that all plants in our experiment were similarly induced. However, a recent study on *B. nigra* highlights the importance of the timing of herbivore attack, making this hypothesis less likely (Rusman *et al*. 2020).

## Conclusion

Our results show that early-season herbivory can affect plant development, the plant-associated herbivore community, and ultimately plant reproduction, and that these effects are highly dependent on the environmental conditions in which they play out. Importantly, effects of early-season herbivory were small and not general across closely related species in similar abiotic and biotic environments suggesting that plant species may be exposed to different levels of natural selection by early-season herbivores through plant- or community-mediated effects on reproduction. However, as plants are often able to compensate and attenuate the effects induced by herbivory, the detection, sign, and strength of effects on seed production are highly dependent on the plant species and environment, indicating that the observed selective potential of early-season herbivory on plant life-history or defence strategies in our study system seems ubiquitously low.

## Acknowledgements

We thank the staff of Unifarm for setting up and maintaining the experimental fields and for processing the seeds. We thank Marcel Dicke and Richard Karban for constructive comments on an earlier version of this manuscript. This project was supported by the European Research Council (ERC) under the European Union’s Horizon 2020 research and innovation program (Grant agreement No. 677139 to EHP) and the NWO RUBICON program (Grant No. 13204 to DM).

## Author contributions

DM and EHP conceived the study and designed the experiment. BK, DM, YZ, and SAZ collected the data. DM and JCD analysed the data. DM and EHP led the writing of the manuscript. All authors provided critical feedback on earlier drafts, contributed substantially to the final version of the manuscript, and gave final approval for publication.

## Data availability

All data and code will be deposited at the *Dryad* repository and will be made publicly available upon publication.

## References

Abdala-Roberts, L., Pérez Niño, B., Moreira, X., Parra-Tabla, V., Grandi, L., Glauser, G. et al. (2019). Effects of early-season insect herbivory on subsequent pathogen infection and ant abundance on wild cotton (Gossypium hirsutum). Journal of Ecology, 107, 1518–1529.

Adhikari, S. & Russell, F.L. (2014). Effects of apical meristem mining on plant fitness, architecture, and flowering phenology in Cirsium altissimum (Asteraceae). American Journal of Botany, 101, 2079–2087.

Agrawal, A.A. (2005). Future directions in the study of induced plant responses to herbivory. Entomologia Experimentalis et Applicata, 115, 97–105.

Ali, J.G. & Agrawal, A.A. (2012). Specialist versus generalist insect herbivores and plant defense. Trends in plant science, 17, 293–302.

Barber, N.A., Adler, L.S., Theis, N., Hazzard, R.V. & Kiers, E.T. (2012). Herbivory reduces plant interactions with above- and belowground antagonists and mutualists. Ecology, 93, 1560–1570.

Bates, D., Mächler, M., Bolker, B. & Walker, S. (2015). Fitting linear mixed-effects models using lme4. Journal of Statistical Software, 67, 1–48.

Behmer, S.T. (2009). Insect herbivore nutrient regulation. Annual Review of Entomology, 54, 165–187.

Boege, K. & Marquis, R.J. (2005). Facing herbivory as you grow up: The ontogeny of resistance in plants. Trends in Ecology & Evolution, 20, 441–448.

Chauta, A., Whitehead, S., Amaya-Marquez, M. & Poveda, K. (2017). Leaf herbivory imposes fitness costs mediated by hummingbird and insect pollinators. PLoS ONE, 12, e0188408.

Cipollini, D., Walters, D. & Voelckel, C. (2014). Costs of resistance in plants: From theory to evidence. In: Insect-Plant Interactions (eds. Voelckel, C & Jander, G). Wiley-Blackwell Oxford, pp. 263–307.

de Bobadilla, M.F., Van Wiechen, R., Gort, G. & Poelman, E.H. (2022). Plasticity in induced resistance to sequential attack by multiple herbivores in Brassica nigra. Oecologia, 198, 11–20.

de Vries, J., Evers, J.B. & Poelman, E.H. (2017). Dynamic plant-plant-herbivore interactions govern plant growth-defence integration. Trends in Plant Science, 22, 329–337.

Dormann, C.F., Gruber, B. & Fründ, J. (2008). Introducing the bipartite package: Analysing ecological networks. Interaction

Douma, J.C. & Shipley, B. (2021). A multigroup extension to piecewise path analysis. Ecosphere, 12, e03502.

Dussourd, D.E. (2017). Behavioral sabotage of plant defenses by insect folivores. Annual Review of Entomology, 62, 15–34.

Edwards, P.J. & Wratten, S.D. (1983). Wound induced defences in plants and their consequences for patterns of insect grazing. Oecologia, 59, 88–93.

Ekholm, A., Tack, A.J.M., Pulkkinen, P. & Roslin, T. (2020). Host plant phenology, insect outbreaks and herbivore communities: The importance of timing. Journal of Animal Ecology, 89, 829–841.

Fox, J. & Weisberg, S. (2018). An R companion to applied regression. Sage Publications.

Garcia, L.C. & Eubanks, M.D. (2019). Overcompensation for insect herbivory: A review and meta-analysis of the evidence. Ecology, 100, e02585.

Garcia, M.B. & Ehrlen, J. (2002). Reproductive effort and herbivory timing in a perennial herb: fitness components at the individual and population levels. Am J Bot, 89, 1295–1302.

Gruntman, M. & Novoplansky, A. (2011). Ontogenetic contingency of tolerance mechanisms in response to apical damage. Ann Bot, 108, 965–973.

Hambäck, P.A., Dahlgren, J.P., Andersson, P., Rabasa, S.G., Bommarco, R. & Ehrlén, J. (2015). Plant trait-mediated interactions between early and late herbivores on common figwort (Scrophularia nodosa) and effects on plant seed set. Écoscience, 18, 375–381.

Han, P., Becker, C., Le Bot, J., Larbat, R., Lavoir, A.V., Desneux, N. et al. (2020). Plant nutrient supply alters the magnitude of indirect interactions between insect herbivores: From foliar chemistry to community dynamics. Journal of Ecology, 108, 1497–1510.

Harrison, X.A., Donaldson, L., Correa-Cano, M.E., Evans, J., Fisher, D.N., Goodwin, C.E.D. et al. (2018). A brief introduction to mixed effects modelling and multi-model inference in ecology. PeerJ, 6, e4794.

Hernandez-Cumplido, J., Glauser, G. & Benrey, B. (2016). Cascading effects of early-season herbivory on late-season herbivores and their parasitoids. Ecology, 97, 1283–1297.

Hoffmeister, M., Wittköpper, N. & Junker, R.R. (2016). Herbivore-induced changes in flower scent and morphology affect the structure of flower–visitor networks but not plant reproduction. Oikos, 125, 1241–1249.

Hothorn, T., Bretz, F. & Westfall, P. (2008). Simultaneous inference in general parametric models. Biom J, 50, 346–363.

Howard, M.M., Kao-Kniffin, J. & Kessler, A. (2020). Shifts in plant-microbe interactions over community succession and their effects on plant resistance to herbivores. New Phytologist, 226, 1144–1157.

Huang, W., Robert, C.A.M., Herve, M.R., Hu, L.F., Bont, Z. & Erb, M. (2017). A mechanism for sequence specificity in plant-mediated interactions between herbivores. New Phytologist, 214, 169–179.

Johnson, J.B. & Omland, K.S. (2004). Model selection in ecology and evolution. Trends in Ecology & Evolution, 19, 101–108.

Kafle, D., Wurst, S. & Dam, N. (2018). Legacy effects of herbivory enhance performance and resistance of progeny plants. Journal of Ecology, 107, 58–68.

Karban, R. (1993). Induced Resistance and Plant Density of a Native Shrub, Gossypium Thurberi, Affect Its Herbivores. Ecology, 74, 1–8.

Karban, R. (2011). The ecology and evolution of induced resistance against herbivores. Functional Ecology, 25, 339–347.

Karban, R. (2019). The ecology and evolution of induced responses to herbivory and how plants perceive risk. Ecological Entomology, 45, 1–9.

Kessler, A. & Halitschke, R. (2007). Specificity and complexity: The impact of herbivore-induced plant responses on arthropod community structure. Current Opinion in Plant Biology, 10, 409–414.

Kindt, R. & Coe, R. (2005). Tree diversity analysis: A manual and software for common statistical methods for ecological and biodiversity studies. World Agroforestry Centre, Nairobi.

Kolb, A., Leimu, R. & Ehrlén, J. (2007). Environmental context influences the outcome of a plant-seed predator interaction. Oikos, 116, 864–872.

Kos, M., Broekgaarden, C., Kabouw, P., Oude Lenferink, K., Poelman, E.H., Vet, L.E.M. et al. (2011). Relative importance of plant-mediated bottom-up and top-down forces on herbivore abundance on Brassica oleracea. Functional Ecology, 25, 1113–1124.

Kostenko, O., van de Voorde, T.F., Mulder, P.P., van der Putten, W.H. & Martijn Bezemer, T. (2012). Legacy effects of aboveground-belowground interactions. Ecology Letters, 15, 813–821.

Lefcheck, J.S. & Freckleton, R. (2015). piecewiseSEM : Piecewise structural equation modelling in R for ecology, evolution, and systematics. Methods in Ecology and Evolution, 7, 573–579.

Legendre, P. & Gallagher, E.D. (2001). Ecologically meaningful transformations for ordination of species data. Oecologia, 129, 271–280.

Lehndal, L. & Agren, J. (2015). Herbivory Differentially Affects Plant Fitness in Three Populations of the Perennial Herb Lythrum salicaria along a Latitudinal Gradient. PLoS One, 10, e0135939.

Lenth, R., Singmann, H., Love, J., Buerkner, P. & Herve, M. (2019). Emmeans: Estimated marginal means, a.k.a. least-squares means. In: R package version 1.1, pp. 1–67.

Li, Y., Stam, J.M., Poelman, E.H., Dicke, M. & Gols, R. (2016). Community structure and abundance of insects in response to early-season aphid infestation in wild cabbage populations. Ecological Entomology, 41, 378–388.

Lill, J.T. & Marquis, R.J. (2003). Ecosystem engineering by caterpillars increases insect herbivore diversity on white oak. Ecology, 84, 682–690.

Lu, J., Robert, C.A., Lou, Y. & Erb, M. (2016). A conserved pattern in plant-mediated interactions between herbivores. Ecology & Evolution, 6, 1032–1040.

Lucas-Barbosa, D., Dicke, M., Kranenburg, T., Aartsma, Y., Beek, T.A., Huigens, M.E. et al. (2016). Endure and call for help: Strategies of black mustard plants to deal with a specialized caterpillar. Functional Ecology, 31, 325–333.

Machado, R.A.R., Arce, C.C.M., McClure, M.A., Baldwin, I.T. & Erb, M. (2018). Aboveground herbivory induced jasmonates disproportionately reduce plant reproductive potential by facilitating root nematode infestation. Plant Cell and Environment, 41, 797–808.

McArt, S.H., Halitschke, R., Salminen, J.P. & Thaler, J.S. (2013). Leaf herbivory increases plant fitness via induced resistance to seed predators. Ecology, 94, 966–975.

Mertens, D., Boege, K., Kessler, A., Koricheva, J., Thaler, J.S., Whiteman, N.K. et al. (2021a). Predictability of biotic stress structures plant defence evolution. Trends in Ecology & Evolution, 36, 444–456.

Mertens, D., Fernández de Bobadilla, M., Rusman, Q., Bloem, J., Douma, J.C. & Poelman, E.H. (2021b). Plant defence to sequential attack is adapted to prevalent herbivores. Nature Plants, 10, 1347–1353.

Mertens, D., Bouwmeester, K. & Poelman, E.H. (2021c). Intraspecific variation in plant-associated herbivore communities is phylogenetically structured in Brassicaceae. Ecol Lett, 24, 2314–2327.

Ochoa-Lopez, S., Villamil, N., Zedillo-Avelleyra, P. & Boege, K. (2015). Plant defence as a complex and changing phenotype throughout ontogeny. Annals of Botany, 116, 797–806.

Ohgushi, T. (2005). Indirect interaction webs: Herbivore-induced effects through trait change in plants. Annual Review of Ecology, Evolution, and Systematics, 36, 81–105.

Oksanen, J., Blanchet, F.G., Kindt, R., Legendre, P., O’Hara, R.B., Simpson, G.L. et al. (2012). Vegan: Community ecology package In: R package version 1.2.

Orrock, J.L., Sih, A., Ferrari, M.C.O., Karban, R., Preisser, E.L., Sheriff, M.J. et al. (2015). Error management in plant allocation to herbivore defense. Trends Ecol. Evol., 30, 441–445.

Pearse, I.S., McMunn, M. & Yang, L.H. (2018). Seasonal assembly of arthropod communities on milkweeds experiencing simulated herbivory. Arthropod-Plant Interactions, 13, 99–108.

Pinheiro, J., Bates, D., DebRoy, S., Sarkar, D. & Team, R.C. (2012). Nlme: Linear and nonlinear mixed effects models. In: R package version 1.1.

Poelman, E.H. & Kessler, A. (2016). Keystone herbivores and the evolution of plant defenses. Trends in Plant Science, 21, 477–485.

R Core Team (2014). R: A language and environment for statistical computing. R Foundation for Statistical Computing Vienna.

Rasmussen, N.L. & Yang, L.H. (2022). Timing of a plant-herbivore interaction alters plant growth and reproduction. Ecology, e3854.

Rubin, I.N., Ellner, S.P., Kessler, A. & Morrell, K.A. (2015). Informed herbivore movement and interplant communication determine the effects of induced resistance in an individual-based model. J Anim Ecol, 84, 1273–1285.

Rusman, Q., Lucas-Barbosa, D., Hassan, K. & Poelman, E.H. (2020). Plant ontogeny determines strength and associated plant fitness consequences of plant-mediated interactions between herbivores and flower visitors. J. Ecol., 108, 1046–1060.

Rusman, Q., Poelman, E.H., Nowrin, F., Polder, G. & Lucas-Barbosa, D. (2019). Floral plasticity: Herbivore-species-specific-induced changes in flower traits with contrasting effects on pollinator visitation. Plant Cell and Environment, 42, 1882–1896.

Santamaria, L. & Rodriguez-Girones, M.A. (2007). Linkage rules for plant-pollinator networks: Trait complementarity or exploitation barriers? PLoS Biology, 5, e31.

Schlinkert, H., Westphal, C., Clough, Y., Grass, I., Helmerichs, J. & Tscharntke, T. (2016). Plant size affects mutualistic and antagonistic interactions and reproductive success across 21 Brassicaceae species. Ecosphere, 7.

Schuman, M.C. & Baldwin, I.T. (2016). The layers of plant responses to insect herbivores. Annual Review of Entomology, 61, 373–394.

Shipley, B. (2009). Confirmatory path analysis in a generalized multilevel context. Ecology, 90, 363–368.

Shipley, B. (2016). Cause and correlation in biology: a user’s guide to path analysis, structural equations and causal inference with R. 2 edn. Cambridge University Press.

Shipley, B. & Douma, J.C. (2020). Generalized AIC and chi-squared statistics for path models consistent with directed acyclic graphs. Ecology, 101, e02960.

Stam, J.M., Dicke, M. & Poelman, E.H. (2018). Order of herbivore arrival on wild cabbage populations influences subsequent arthropod community development. Oikos, 127, 1482–1493.

Stam, J.M., Kos, M., Dicke, M. & Poelman, E.H. (2019). Cross-seasonal legacy effects of arthropod community on plant fitness in perennial plants. Journal of Ecology, 107, 2451–2463.

Strauss, S.Y. & Agrawal, A.A. (1999). The ecology and evolution of plant tolerance to herbivory. Trends in Ecology & Evolution, 14, 179–185.

Tonkin, J.D., Bogan, M.T., Bonada, N., Rios-Touma, B. & Lytle, D.A. (2017). Seasonality and predictability shape temporal species diversity. Ecology, 98, 1201–1216.

Visakorpi, K., Riutta, T., Martínez-Bauer, A.E., Salminen, J.P. & Gripenberg, S. (2019). Insect community structure covaries with host plant chemistry but is not affected by prior herbivory. Ecology, 100, e02739.

Viswanathan, D.V., Narwani, A.J.T. & Thaler, J.S. (2005). Specificity in induced plant responses shapes patterns of herbivore occurrence on Solanum dulcamara. Ecology, 86, 886–896.

West, N.M. & Louda, S.M. (2018). Cumulative herbivory outpaces compensation for early floral damage on a monocarpic perennial thistle. Oecologia, 186, 495–506.

Wickham, H. (2009). ggplot2: elegant graphics for data analysis. Springer Science & Business Media, New York.

Williams, I.H. (2010). The Major Insect Pests of Oilseed Rape in Europe and Their Management: An Overview. In: Biocontrol-Based Integrated Management of Oilseed Rape Pests (ed. Williams, IH). Springer Dordrecht, pp. 1–43.

Wold, E.N. & Marquis, R.J. (1997). Induced Defense in White Oak: Effects on Herbivores and Consequences for the Plant. Ecology, 78, 1356–1369.

Wood, S., Scheipl, F. & Wood, M.S. (2017). Package ‘gamm4’. In: American Statistical Association, p. 339.

Wood, S.N. (2011). Fast stable restricted maximum likelihood and marginal likelihood estimation of semiparametric generalized linear models. Journal of the Royal Statistical Society: Series B (Statistical Methodology), 73, 3–36.

Wright, S.D. & McConnaughay, K.D.M. (2002). Interpreting phenotypic plasticity: The importance of ontogeny. Plant Species Biology, 17, 119–131.

Wurst, S., Ohgushi, T. & Allen, E. (2015). Do plant- and soil-mediated legacy effects impact future biotic interactions? Functional Ecology, 29, 1373–1382.

Zuur, A., Ieno, E.N., Walker, N., Saveliev, A.A. & Smith, G.M. (2009). Mixed Effects Models and Extensions in Ecology with R. Springer Science & Business Media, New York.

