## Supplemental tables and figures for "Priority effects in herbivore communities vary in effect on plant development and reproduction in four Brassicaceae plant species"

This document contains:

- Supplemental tables S1 – S10
- Supplemental figures S1 – S8

**Table S1.** Plant species used in our experiment, detailing the number of plants included per treatment in the two field seasons as used in the statistical analysis. NF details the number of plants in the open-field experiment, NE details the number of plants in the community exclosure experiment. The subscript indicates the year of the field season.

| Species name | Treatment | $NF_{2017}$ | $NE_{2017}$ | $NF_{2018}$ | $NE_{2018}$ |
|---|---|---|---|---|---|
| *Brassica nigra* | *Myzus persicae* | 39 | 8 | 40 | 15 |
| | *Pieris rapae* | 32 | 8 | 40 | 12 |
| | Untreated | 34 | 9 | 40 | 11 |
| *Raphanus raphanistrum* | *Myzus persicae* | 39 | 9 | 40 | 15 |
| | *Pieris rapae* | 35 | 4 | 39 | 15 |
| | Untreated | 38 | 9 | 40 | 17 |
| *Sinapis arvensis* | *Myzus persicae* | 25 | 7 | 40 | 16 |
| | *Pieris rapae* | 29 | 4 | 39 | 14 |
| | Untreated | 28 | 8 | 35 | 14 |
| *Rapistrum rugosum* | *Myzus persicae* | 38 | 6 | 39 | 9 |
| | *Pieris rapae* | 29 | 7 | 38 | 9 |
| | Untreated | 37 | 8 | 36 | 7 |

**Table S2**. Herbivore insect species observed in our experiment. Species annotated with an asterisk were included in our data as morphospecies. The species for which we manipulated presence and abundance as treatment in our experiment are indicated in bold and were excluded from the data.

| **Calling name** | | **Species name** | **Order** | **Family** |
|---|---|---|---|---|
| Weevils | * | - | Coleoptera | - |
| Leaf beetle | * | - | Coleoptera | Chrysomelidae |
| Red flea beetle | | *Neocrepidodera transversa* | Coleoptera | Chrysomelidae |
| Mustard beetle | | *Phaedon cochleariae* | Coleoptera | Chrysomelidae |
| Turnip flea beetle | | *Phyllotreta atra* | Coleoptera | Chrysomelidae |
| Small striped flea beetle | | *Phyllotreta undulata* | Coleoptera | Chrysomelidae |
| Pollen beetle | | *Brassicogethes aeneus* | Coleoptera | Nitidulidae |
| Collembola | * | - | Collembola | - |
| Leaf mining flies | * | - | Diptera | - |
| Cicades | * | - | Hemiptera | - |
| Black bean aphid | | *Aphis fabae* | Hemiptera | Aphididae |
| Cabbage aphid | | *Brevicoryne brassicae* | Hemiptera | Aphididae |
| Mustard aphid | | *Lipaphis erysimi* | Hemiptera | Aphididae |
| **Green peach aphid** | | ***Myzus persicae*** | Hemiptera | Aphididae |
| Tobacco aphid | | *Myzus persicae* sub. *nicotianae* | Hemiptera | Aphididae |
| Harlequin bug | | *Murgantia histrionica* | Hemiptera | Pentatomidae |
| Lygus bug | * | *Lygus spp.* | Hemiptera | Miridae |
| Turnip sawfly | | *Athalia rosae* | Hymenoptera | Tenthredinidae |
| Loopers | * | - | Lepidoptera | - |
| Marbled yellow pearl | | *Evergestis extimalis* | Lepidoptera | Crambidae |
| Silver Y | | *Autographa gamma* | Lepidoptera | Noctuidae |
| Cabbage moth | | *Mamestra brassicae* | Lepidoptera | Noctuidae |
| Large cabbage white | | *Pieris brassicae* | Lepidoptera | Pieridae |
| **Small cabbage white** | | ***Pieris rapae*** | Lepidoptera | Pieridae |
| Diamondback moth | | *Plutella xylostella* | Lepidoptera | Yponomeutidae |
| Thrips | * | - | Thysanoptera | - |

**Table S3.** Overview of differently formulated path models, the degrees of freedom, and their fit expressed by AIC, and Fisher's C global goodness of fit statistic with the associated p-value. A p > 0.05 indicates that the data are sufficiently well represented by the path model. Each model places different constraints on the mean estimation of variables and path coefficients. **A:** Path coefficients and estimation of variables constrained to be equal across all plant species and the two years. **B:** Path coefficients are constrained, estimations of the mean of variables in the model are allowed to vary among plant species and the two years in an additive way. **C:** Path coefficients are constrained, estimations of the mean of variables are allowed to vary among plant species and the two years in an interactive way. **D:** Path coefficients are constrained, and estimations of variables in the model are allowed to vary among plant species and the two years in an interactive way, where plant species are included as a crossed random effect (i.e. assumed to come from a normal distributed representation of Brassicaceae plant species). **E:** Path coefficients are constrained to be equal among plant species but can vary across the two years. Estimations of the mean of variables in the model are allowed to vary among plant species. **F:** Path coefficients are constrained to be equal among the two years but can vary across the different plant species. Estimations of the mean of variables in the model are allowed to vary among the two years. **G:** Path coefficients as well as mean estimations of variables in the model can vary among plant species and the two years. Note that the models using approach E, F, and G are fitted to subsets of the data according to the year and plant species, making models not directly comparable in terms of fit.

| Model | Year | Plant Species | df | AIC | Fisher's C | p-value |
|---|---|---|---|---|---|---|
| **A** | 2017 & 2018 | All | 74 | 604.957 | 472.957 | < 0.0001 |
| **B** | 2017 & 2018 | All | 74 | 717.774 | 437.774 | < 0.0001 |
| **C** | 2017 & 2018 | All | 74 | 866.151 | 478.151 | < 0.0001 |
| **D** | 2017 & 2018 | All | 74 | 706.315 | 510.315 | < 0.0001 |
| **E** | 2017 | All | 74 | 475.485 | 229.485 | < 0.0001 |
| | 2018 | All | 74 | 595.947 | 349.947 | < 0.0001 |
| **F** | 2017 & 2018 | *Brassica nigra* | 74 | 373.668 | 207.668 | < 0.0001 |
| | 2017 & 2018 | *Raphanus raphanistrum* | 74 | 329.842 | 163.842 | < 0.0001 |
| | 2017 & 2018 | *Sinapis arvensis* | 74 | 355.712 | 189.712 | < 0.0001 |
| | 2017 & 2018 | *Rapistrum rugosum* | 74 | 409.793 | 243.793 | < 0.0001 |
| **G** | 2017 | *Brassica nigra* | 74 | 303.737 | 171.737 | < 0.0001 |
| | 2017 | *Raphanus raphanistrum* | 74 | 246.239 | 114.239 | < 0.0001 |
| | 2017 | *Sinapis arvensis* | 74 | 245.879 | 113.879 | < 0.0001 |
| | 2017 | *Rapistrum rugosum* | 74 | 287.164 | 155.164 | < 0.0001 |
| | 2018 | *Brassica nigra* | 74 | 274.903 | 142.903 | < 0.0001 |
| | 2018 | *Raphanus raphanistrum* | 74 | 267.533 | 135.533 | < 0.0001 |
| | 2018 | *Sinapis arvensis* | 74 | 314.759 | 182.759 | < 0.0001 |
| | 2018 | *Rapistrum rugosum* | 74 | 299.032 | 167.032 | < 0.0001 |

**Table S4.** Overview of the fit and structure of path models after optimization when starting from path models with different constraints on the effects of plant species and year on the mean estimated for variables in the path models. Models are annotated with their degrees of freedom, and their fit expressed by AIC and Fisher's C global goodness of fit statistic with the associated p-value. A p-value > 0.05 indicates that the data are sufficiently well represented by the path model. The table further presents the number of paths included in the model, the number of correlated error structures, and the number of independence claims, along with the number of paths, correlated error structures, and claims which were found to be significant. **A:** Path coefficients and estimation of variables constrained to be equal across all plant species and the two years. **B:** Path coefficients are constrained, estimations of the mean of variables in the model are allowed to vary among plant species and the two years in an additive way. **C:** Path coefficients are constrained, estimations of the mean of variables are allowed to vary among plant species and the two years in an interactive way. **D:** Path coefficients are constrained, and estimations of variables in the model are allowed to vary among plant species and the two years in an interactive way, where plant species are included as a crossed random effect (i.e. plant species are assumed to originate from a normal distributed representation of Brassicaceae plant species).

| Model | df | AIC | Fisher's C | p-value | Paths | Significant paths | Error structures | Significant error structures | Independence Claims | Significant independence claims |
|---|---|---|---|---|---|---|---|---|---|---|
| **A** | 48 | 157.271 | 43.271 | 0.6670 | 25 | 21 | 4 | 4 | 24 | 0 |
| **B** | 74 | 238.627 | 73.487 | 0.4950 | 22 | 16 | 6 | 5 | 37 | 0 |
| **C** | 82 | 458.442 | 84.442 | 0.4050 | 19 | 17 | 5 | 4 | 41 | 0 |
| **D** | 78 | 345.487 | 58.627 | 0.9500 | 20 | 17 | 6 | 5 | 39 | 0 |

**Table S5.** Overview of path coefficients and their associated standard error (Estimate ± SE) and p-value for each of the causal relations retained in the optimised descriptive piecewise path model. Estimations were obtained by fitting the model to subsets of the full data for each of the two years, the four plant species, or each plant species by year combination. The p-values of path coefficients which significantly differ from zero ($p < 0.05$) are indicated in bold. The volume of plants and the number of reproductive branches were square-root-transformed, and the abundances of folivores, florivores and seed predators were transformed using a log(x+1) transformation.

| | | | 2017 & 2018 | | 2017 | | 2018 | |
|---|---|---|---|---|---|---|---|---|
| **Response** | **Predictor** | **Plant species** | **Estimate ± SE** | **p-value** | **Estimate ± SE** | **p-value** | **Estimate ± SE** | **p-value** |
| Early-season herbivore richness | Early-season number of leaves | All | 0.19 ± 0.04 | **< 0.0001** | 0.12 ± 0.10 | **0.0269** | 0.19 ± 0.06 | **0.0005** |
| | | *Brassica nigra* | 0.34 ± 0.10 | **< 0.0001** | 0.23 ± 0.21 | **0.0271** | 0.30 ± 0.15 | **0.0014** |
| | | *Raphanus raphanistrum* | 0.12 ± 0.06 | 0.1389 | 0.06 ± 0.15 | 0.5374 | 0.01 ± 0.10 | 0.9056 |
| | | *Rapistrum rugosum* | 0.12 ± 0.06 | 0.1389 | 0.06 ± 0.15 | 0.5374 | 0.01 ± 0.10 | 0.9056 |
| | | *Sinapis arvensis* | 0.08 ± 0.12 | 0.2958 | 0.01 ± 0.21 | 0.9463 | 0.14 ± 0.16 | 0.1361 |
| Mid-season number of leaves | Early-season number of leaves | All | 0.35 ± 0.04 | **< 0.0001** | 0.35 ± 0.07 | **< 0.0001** | 0.26 ± 0.06 | **< 0.0001** |
| | | *Brassica nigra* | 0.26 ± 0.09 | **0.0001** | 0.36 ± 0.16 | **0.0003** | 0.14 ± 0.12 | 0.1223 |
| | | *Raphanus raphanistrum* | 0.33 ± 0.06 | **0.0012** | 0.33 ± 0.13 | **0.0013** | 0.18 ± 0.08 | 0.0766 |
| | | *Rapistrum rugosum* | 0.33 ± 0.06 | **0.0012** | 0.33 ± 0.13 | **0.0013** | 0.18 ± 0.08 | 0.0766 |
| | | *Sinapis arvensis* | 0.35 ± 0.08 | **0.0000** | 0.35 ± 0.13 | **0.0034** | 0.31 ± 0.10 | **0.0028** |
| Mid-season number of leaves | Early-season volume | All | 0.07 ± 0.05 | 0.1672 | 0.15 ± 0.07 | **0.0058** | -0.04 ± 0.07 | 0.5892 |
| | | *Brassica nigra* | 0.38 ± 0.12 | **< 0.0001** | 0.17 ± 0.23 | 0.0649 | 0.27 ± 0.17 | **0.0051** |
| | | *Raphanus raphanistrum* | 0.25 ± 0.07 | **0.0151** | 0.18 ± 0.11 | 0.0611 | 0.07 ± 0.11 | 0.5092 |
| | | *Rapistrum rugosum* | 0.25 ± 0.07 | **0.0151** | 0.18 ± 0.11 | 0.0611 | 0.07 ± 0.11 | 0.5092 |
| | | *Sinapis arvensis* | -0.07 ± 0.07 | 0.4452 | 0.14 ± 0.12 | 0.2900 | -0.11 ± 0.09 | 0.2782 |
| Mid-season number of leaves | Early-season herbivore abundance | All | 0.09 ± 0.03 | **0.0007** | 0.07 ± 0.02 | 0.0524 | 0.13 ± 0.05 | **0.0026** |
| | | *Brassica nigra* | 0.06 ± 0.05 | 0.2155 | 0.10 ± 0.05 | 0.1727 | 0.07 ± 0.09 | 0.4784 |
| | | *Raphanus raphanistrum* | 0.13 ± 0.05 | **0.0204** | 0.04 ± 0.05 | 0.5161 | 0.23 ± 0.08 | **0.0145** |
| | | *Rapistrum rugosum* | 0.13 ± 0.05 | **0.0204** | 0.04 ± 0.05 | 0.5161 | 0.23 ± 0.08 | **0.0145** |
| | | *Sinapis arvensis* | 0.11 ± 0.04 | 0.1146 | 0.17 ± 0.05 | 0.0589 | -0.09 ± 0.07 | 0.3841 |
| Mid-season volume | Early-season volume | All | 0.05 ± 0.04 | 0.2889 | 0.04 ± 0.11 | 0.4540 | 0.29 ± 0.04 | **< 0.0001** |
| | | *Brassica nigra* | 0.19 ± 0.12 | **0.0238** | 0.41 ± 0.33 | **< 0.0001** | 0.51 ± 0.12 | **< 0.0001** |
| | | *Raphanus raphanistrum* | -0.27 ± 0.04 | **0.0003** | 0.02 ± 0.14 | 0.8615 | 0.24 ± 0.06 | **0.0086** |
| | | *Rapistrum rugosum* | -0.27 ± 0.04 | **0.0003** | 0.02 ± 0.14 | 0.8615 | 0.24 ± 0.06 | **0.0086** |
| | | *Sinapis arvensis* | -0.05 ± 0.08 | 0.5116 | -0.31 ± 0.14 | **0.0066** | 0.35 ± 0.06 | **0.0001** |

**Table S5. Continued**

| | | | 2017 & 2018 | | 2017 | | 2018 | |
|---|---|---|---|---|---|---|---|---|
| **Response** | **Predictor** | **Plant species** | **Estimate ± SE** | **p-value** | **Estimate ± SE** | **p-value** | **Estimate ± SE** | **p-value** |
| Mid-season volume | Early-season herbivore abundance | All | 0.13 ± 0.03 | **< 0.0001** | 0.09 ± 0.05 | 0.0642 | 0.21 ± 0.03 | **< 0.0001** |
| | | *Brassica nigra* | 0.09 ± 0.06 | 0.1848 | 0.05 ± 0.09 | 0.6084 | 0.00 ± 0.06 | 0.9828 |
| | | *Raphanus raphanistrum* | 0.13 ± 0.05 | 0.0521 | 0.11 ± 0.08 | 0.2632 | 0.24 ± 0.05 | **0.0125** |
| | | *Rapistrum rugosum* | 0.13 ± 0.05 | 0.0521 | 0.11 ± 0.08 | 0.2632 | 0.24 ± 0.05 | **0.0125** |
| | | *Sinapis arvensis* | 0.05 ± 0.05 | 0.4578 | 0.03 ± 0.08 | 0.7665 | 0.11 ± 0.04 | 0.2252 |
| Mid-season herbivore richness | Early-season volume | All | -0.09 ± 0.04 | **0.0249** | 0.02 ± 0.12 | 0.8001 | -0.08 ± 0.05 | 0.1714 |
| | | *Brassica nigra* | -0.08 ± 0.13 | 0.3447 | -0.15 ± 0.40 | 0.2080 | 0.00 ± 0.16 | 0.9711 |
| | | *Raphanus raphanistrum* | -0.25 ± 0.07 | **0.0031** | 0.00 ± 0.22 | 0.9850 | -0.14 ± 0.11 | 0.1599 |
| | | *Rapistrum rugosum* | -0.25 ± 0.07 | **0.0031** | 0.00 ± 0.22 | 0.9850 | -0.14 ± 0.11 | 0.1599 |
| | | *Sinapis arvensis* | 0.03 ± 0.12 | 0.7370 | 0.03 ± 0.25 | 0.8480 | 0.02 ± 0.15 | 0.8089 |
| Mid-season herbivore richness | Early-season herbivore abundance | All | 0.12 ± 0.03 | **0.0004** | 0.08 ± 0.05 | 0.1194 | 0.15 ± 0.05 | **0.0020** |
| | | *Brassica nigra* | 0.08 ± 0.07 | 0.2330 | 0.06 ± 0.10 | 0.5721 | 0.10 ± 0.09 | 0.2683 |
| | | *Raphanus raphanistrum* | 0.17 ± 0.07 | **0.0125** | 0.12 ± 0.11 | 0.2317 | 0.17 ± 0.09 | 0.0802 |
| | | *Rapistrum rugosum* | 0.17 ± 0.07 | **0.0125** | 0.12 ± 0.11 | 0.2317 | 0.17 ± 0.09 | 0.0802 |
| | | *Sinapis arvensis* | 0.14 ± 0.08 | 0.0576 | 0.12 ± 0.12 | 0.2937 | 0.12 ± 0.11 | 0.2329 |
| Mid-season herbivore richness | Mid-season number of leaves | All | -0.11 ± 0.04 | **0.0027** | -0.05 ± 0.09 | 0.4491 | -0.11 ± 0.04 | **0.0337** |
| | | *Brassica nigra* | -0.15 ± 0.08 | 0.0634 | 0.06 ± 0.17 | 0.6136 | -0.21 ± 0.09 | **0.0233** |
| | | *Raphanus raphanistrum* | -0.01 ± 0.08 | 0.8879 | -0.01 ± 0.19 | 0.9198 | -0.03 ± 0.10 | 0.7767 |
| | | *Rapistrum rugosum* | -0.01 ± 0.08 | 0.8879 | -0.01 ± 0.19 | 0.9198 | -0.03 ± 0.10 | 0.7767 |
| | | *Sinapis arvensis* | 0.03 ± 0.15 | 0.7149 | 0.15 ± 0.32 | 0.3362 | -0.03 ± 0.17 | 0.7499 |

**Table S5. Continued**

| | | | 2017 & 2018 | | 2017 | | 2018 | |
|---|---|---|---|---|---|---|---|---|
| **Response** | **Predictor** | **Plant species** | **Estimate ± SE** | **p-value** | **Estimate ± SE** | **p-value** | **Estimate ± SE** | **p-value** |
| Mid-season herbivore abundance | Early-season herbivore abundance | All | 0.10 ± 0.03 | **0.0024** | 0.06 ± 0.05 | 0.2554 | 0.14 ± 0.05 | **0.0044** |
| | | *Brassica nigra* | 0.05 ± 0.07 | 0.4872 | 0.02 ± 0.10 | 0.8550 | 0.07 ± 0.10 | 0.4762 |
| | | *Raphanus raphanistrum* | 0.19 ± 0.07 | **0.0056** | 0.19 ± 0.09 | **0.0450** | 0.20 ± 0.10 | **0.0447** |
| | | *Rapistrum rugosum* | 0.19 ± 0.07 | **0.0056** | 0.19 ± 0.09 | **0.0450** | 0.20 ± 0.10 | **0.0447** |
| | | *Sinapis arvensis* | 0.07 ± 0.08 | 0.3132 | 0.07 ± 0.10 | 0.5712 | 0.10 ± 0.13 | 0.3027 |
| Mid-season herbivore abundance | Mid-season number of leaves | All | -0.10 ± 0.04 | **0.0050** | -0.01 ± 0.08 | 0.9171 | -0.12 ± 0.05 | **0.0150** |
| | | *Brassica nigra* | -0.16 ± 0.07 | **0.0410** | 0.03 ± 0.15 | 0.7676 | -0.23 ± 0.10 | **0.0153** |
| | | *Raphanus raphanistrum* | -0.07 ± 0.08 | 0.2822 | 0.07 ± 0.13 | 0.4837 | -0.07 ± 0.11 | 0.4307 |
| | | *Rapistrum rugosum* | -0.07 ± 0.08 | 0.2822 | 0.07 ± 0.13 | 0.4837 | -0.07 ± 0.11 | 0.4307 |
| | | *Sinapis arvensis* | 0.03 ± 0.14 | 0.6523 | -0.02 ± 0.20 | 0.8756 | 0.07 ± 0.19 | 0.4672 |
| Reproductive branches | Early-season number of leaves | All | 0.09 ± 0.04 | **0.0180** | 0.03 ± 0.11 | 0.5420 | 0.18 ± 0.03 | **0.0001** |
| | | *Brassica nigra* | -0.07 ± 0.11 | 0.2947 | 0.07 ± 0.25 | 0.4586 | -0.05 ± 0.11 | 0.5509 |
| | | *Raphanus raphanistrum* | 0.08 ± 0.05 | 0.4140 | 0.00 ± 0.14 | 0.9794 | 0.24 ± 0.03 | **0.0058** |
| | | *Rapistrum rugosum* | 0.08 ± 0.05 | 0.4140 | 0.00 ± 0.14 | 0.9794 | 0.24 ± 0.03 | **0.0058** |
| | | *Sinapis arvensis* | -0.01 ± 0.04 | 0.8882 | -0.04 ± 0.07 | 0.7072 | 0.01 ± 0.04 | 0.8639 |
| Reproductive branches | Early-season volume | All | -0.12 ± 0.04 | **0.0029** | -0.16 ± 0.11 | **0.0042** | -0.05 ± 0.04 | 0.3794 |
| | | *Brassica nigra* | 0.05 ± 0.15 | 0.4824 | 0.08 ± 0.37 | 0.3968 | 0.18 ± 0.17 | 0.0545 |
| | | *Raphanus raphanistrum* | -0.32 ± 0.05 | **0.0019** | -0.14 ± 0.13 | 0.2405 | 0.11 ± 0.05 | 0.1926 |
| | | *Rapistrum rugosum* | -0.32 ± 0.05 | **0.0019** | -0.14 ± 0.13 | 0.2405 | 0.11 ± 0.05 | 0.1926 |
| | | *Sinapis arvensis* | -0.06 ± 0.03 | 0.4045 | -0.20 ± 0.06 | 0.1106 | 0.05 ± 0.04 | 0.5469 |

**Table S5. Continued**

| Response | Predictor | Plant species | 2017 & 2018 | | 2017 | | 2018 | |
|---|---|---|---|---|---|---|---|---|
| | | | Estimate ± SE | p-value | Estimate ± SE | p-value | Estimate ± SE | p-value |
| Reproductive branches | Mid-season number of leaves | All | 0.03 ± 0.03 | 0.3034 | 0.14 ± 0.08 | **0.0098** | 0.10 ± 0.03 | **0.0432** |
| | | *Brassica nigra* | -0.03 ± 0.08 | 0.6206 | 0.06 ± 0.15 | 0.5472 | -0.10 ± 0.10 | 0.3161 |
| | | *Raphanus raphanistrum* | 0.07 ± 0.05 | 0.3294 | 0.14 ± 0.10 | 0.2139 | 0.28 ± 0.04 | **0.0014** |
| | | *Rapistrum rugosum* | 0.07 ± 0.05 | 0.3294 | 0.14 ± 0.10 | 0.2139 | 0.28 ± 0.04 | **0.0014** |
| | | *Sinapis arvensis* | 0.42 ± 0.03 | **< 0.0001** | 0.36 ± 0.06 | **0.0013** | 0.56 ± 0.04 | **< 0.0001** |
| Reproductive branches | Mid-season herbivore richness | All | -0.05 ± 0.02 | **0.0158** | -0.07 ± 0.03 | 0.0345 | -0.02 ± 0.02 | 0.5614 |
| | | *Brassica nigra* | -0.03 ± 0.06 | 0.5353 | -0.02 ± 0.08 | 0.8217 | -0.05 ± 0.08 | 0.5232 |
| | | *Raphanus raphanistrum* | -0.15 ± 0.03 | **0.0055** | -0.18 ± 0.05 | **0.0283** | -0.11 ± 0.04 | 0.1587 |
| | | *Rapistrum rugosum* | -0.15 ± 0.03 | **0.0055** | -0.18 ± 0.05 | **0.0283** | -0.11 ± 0.04 | 0.1587 |
| | | *Sinapis arvensis* | -0.05 ± 0.01 | 0.3124 | 0.05 ± 0.02 | 0.4889 | -0.14 ± 0.02 | 0.0628 |
| Reproductive branches | Mid-season volume | All | 0.42 ± 0.03 | **< 0.0001** | 0.34 ± 0.04 | **< 0.0001** | 0.36 ± 0.05 | **< 0.0001** |
| | | *Brassica nigra* | 0.24 ± 0.07 | **< 0.0001** | 0.07 ± 0.09 | 0.3930 | 0.28 ± 0.12 | **0.0018** |
| | | *Raphanus raphanistrum* | 0.45 ± 0.05 | **< 0.0001** | 0.46 ± 0.06 | **< 0.0001** | 0.08 ± 0.07 | 0.3657 |
| | | *Rapistrum rugosum* | 0.45 ± 0.05 | **< 0.0001** | 0.46 ± 0.06 | **< 0.0001** | 0.08 ± 0.07 | 0.3657 |
| | | *Sinapis arvensis* | 0.42 ± 0.03 | **< 0.0001** | 0.42 ± 0.04 | **< 0.0001** | 0.16 ± 0.06 | 0.1005 |
| Florivore & seed predator abundance | Early-season number of leaves | All | 0.16 ± 0.05 | **0.0010** | -0.04 ± 0.12 | 0.5634 | 0.07 ± 0.05 | 0.2524 |
| | | *Brassica nigra* | 0.11 ± 0.09 | 0.2035 | 0.12 ± 0.20 | 0.3564 | 0.03 ± 0.10 | 0.7784 |
| | | *Raphanus raphanistrum* | 0.19 ± 0.08 | 0.0622 | -0.09 ± 0.22 | 0.5042 | 0.07 ± 0.08 | 0.4860 |
| | | *Rapistrum rugosum* | 0.19 ± 0.08 | 0.0622 | -0.09 ± 0.22 | 0.5042 | 0.07 ± 0.08 | 0.4860 |
| | | *Sinapis arvensis* | -0.03 ± 0.15 | 0.7399 | -0.14 ± 0.28 | 0.4251 | -0.06 ± 0.15 | 0.5503 |

**Table S5. Continued**

| | | | 2017 & 2018 | | 2017 | | 2018 | |
|---|---|---|---|---|---|---|---|---|
| **Response** | **Predictor** | **Plant species** | **Estimate ± SE** | **p-value** | **Estimate ± SE** | **p-value** | **Estimate ± SE** | **p-value** |
| Florivore & seed predator abundance | Early-season volume | All | 0.12 ± 0.05 | **0.0241** | 0.02 ± 0.11 | 0.8217 | 0.02 ± 0.05 | 0.7596 |
| | | *Brassica nigra* | 0.13 ± 0.12 | 0.1523 | -0.33 ± 0.30 | **0.0109** | 0.09 ± 0.14 | 0.3720 |
| | | *Raphanus raphanistrum* | 0.36 ± 0.09 | **0.0010** | 0.23 ± 0.2 | 0.1003 | 0.12 ± 0.11 | 0.2574 |
| | | *Rapistrum rugosum* | 0.36 ± 0.09 | **0.0010** | 0.23 ± 0.2 | 0.1003 | 0.12 ± 0.11 | 0.2574 |
| | | *Sinapis arvensis* | 0.21 ± 0.13 | **0.0168** | 0.21 ± 0.23 | 0.2333 | 0.03 ± 0.13 | 0.7573 |
| Florivore & seed predator abundance | Mid-season herbivore richness | All | -0.11 ± 0.05 | **0.0306** | 0.01 ± 0.07 | 0.9433 | -0.15 ± 0.07 | 0.0600 |
| | | *Brassica nigra* | -0.08 ± 0.09 | 0.4270 | 0.03 ± 0.11 | 0.8436 | -0.21 ± 0.14 | 0.2284 |
| | | *Raphanus raphanistrum* | 0.03 ± 0.10 | 0.7960 | 0.22 ± 0.14 | 0.2233 | -0.11 ± 0.15 | 0.4993 |
| | | *Rapistrum rugosum* | 0.03 ± 0.10 | 0.7960 | 0.22 ± 0.14 | 0.2233 | -0.11 ± 0.15 | 0.4993 |
| | | *Sinapis arvensis* | -0.06 ± 0.09 | 0.5480 | 0.06 ± 0.15 | 0.7556 | -0.03 ± 0.10 | 0.8113 |
| Florivore & seed predator abundance | Mid-season herbivore abundance | All | 0.12 ± 0.05 | **0.0213** | 0.06 ± 0.07 | 0.4940 | 0.17 ± 0.06 | **0.0293** |
| | | *Brassica nigra* | -0.01 ± 0.09 | 0.9130 | -0.06 ± 0.11 | 0.6959 | 0.03 ± 0.13 | 0.8581 |
| | | *Raphanus raphanistrum* | -0.07 ± 0.10 | 0.4512 | -0.19 ± 0.16 | 0.2712 | 0.06 ± 0.13 | 0.7261 |
| | | *Rapistrum rugosum* | -0.07 ± 0.10 | 0.4512 | -0.19 ± 0.16 | 0.2712 | 0.06 ± 0.13 | 0.7261 |
| | | *Sinapis arvensis* | 0.23 ± 0.09 | **0.0164** | -0.02 ± 0.18 | 0.9296 | 0.42 ± 0.09 | **0.0018** |
| Florivore & seed predator abundance | Reproductive branches | All | -0.11 ± 0.04 | **0.0021** | 0.06 ± 0.04 | 0.2987 | -0.07 ± 0.06 | 0.1907 |
| | | *Brassica nigra* | -0.23 ± 0.05 | **0.0025** | 0.03 ± 0.07 | 0.7913 | -0.19 ± 0.08 | 0.0714 |
| | | *Raphanus raphanistrum* | -0.06 ± 0.10 | 0.3599 | 0.05 ± 0.13 | 0.5847 | -0.02 ± 0.21 | 0.8293 |
| | | *Rapistrum rugosum* | -0.06 ± 0.10 | 0.3599 | 0.05 ± 0.13 | 0.5847 | -0.02 ± 0.21 | 0.8293 |
| | | *Sinapis arvensis* | 0.06 ± 0.23 | 0.3310 | 0.08 ± 0.30 | 0.4750 | 0.24 ± 0.27 | **0.0109** |

**Table S5. Continued**

| Response | Predictor | Plant species | 2017 & 2018 | | 2017 | | 2018 | |
|---|---|---|---|---|---|---|---|---|
| | | | Estimate ± SE | p-value | Estimate ± SE | p-value | Estimate ± SE | p-value |
| Seed set | Early-season volume | All | -0.08 ± 0.04 | **0.0476** | 0.03 ± 0.11 | 0.5971 | -0.14 ± 0.03 | **0.0009** |
| | | *Brassica nigra* | -0.08 ± 0.14 | 0.3278 | -0.09 ± 0.48 | 0.4667 | 0.06 ± 0.17 | 0.5054 |
| | | *Raphanus raphanistrum* | 0.18 ± 0.03 | **0.0366** | 0.12 ± 0.09 | 0.2589 | -0.06 ± 0.04 | 0.4976 |
| | | *Rapistrum rugosum* | 0.18 ± 0.03 | **0.0366** | 0.12 ± 0.09 | 0.2589 | -0.06 ± 0.04 | 0.4976 |
| | | *Sinapis arvensis* | -0.16 ± 0.05 | **0.0028** | -0.01 ± 0.14 | 0.9443 | -0.08 ± 0.05 | 0.2241 |
| Seed set | Mid-season number of leaves | All | 0.19 ± 0.04 | **< 0.0001** | 0.09 ± 0.09 | 0.1051 | 0.33 ± 0.04 | **< 0.0001** |
| | | *Brassica nigra* | 0.16 ± 0.09 | 0.0508 | 0.22 ± 0.18 | 0.0598 | 0.17 ± 0.11 | 0.1008 |
| | | *Raphanus raphanistrum* | 0.10 ± 0.04 | 0.1982 | -0.02 ± 0.08 | 0.8601 | 0.09 ± 0.05 | 0.4104 |
| | | *Rapistrum rugosum* | 0.10 ± 0.04 | 0.1982 | -0.02 ± 0.08 | 0.8601 | 0.09 ± 0.05 | 0.4104 |
| | | *Sinapis arvensis* | 0.09 ± 0.07 | 0.1264 | 0.10 ± 0.17 | 0.4347 | 0.24 ± 0.07 | **0.0096** |
| Seed set | Mid-season volume | All | 0.39 ± 0.03 | **< 0.0001** | 0.38 ± 0.05 | **< 0.0001** | 0.36 ± 0.05 | **< 0.0001** |
| | | *Brassica nigra* | 0.32 ± 0.08 | **< 0.0001** | 0.31 ± 0.12 | **0.0038** | 0.20 ± 0.15 | 0.0811 |
| | | *Raphanus raphanistrum* | 0.21 ± 0.04 | **0.0065** | 0.14 ± 0.06 | 0.1614 | 0.32 ± 0.07 | **0.0014** |
| | | *Rapistrum rugosum* | 0.21 ± 0.04 | **0.0065** | 0.14 ± 0.06 | 0.1614 | 0.32 ± 0.07 | **0.0014** |
| | | *Sinapis arvensis* | 0.27 ± 0.05 | **< 0.0001** | 0.21 ± 0.09 | **0.0349** | -0.01 ± 0.07 | 0.8907 |
| Seed set | Reproductive branches | All | 0.05 ± 0.04 | 0.1657 | 0.13 ± 0.05 | **0.0089** | -0.09 ± 0.05 | **0.0416** |
| | | *Brassica nigra* | 0.01 ± 0.07 | 0.8666 | 0.08 ± 0.11 | 0.4717 | -0.13 ± 0.10 | 0.1790 |
| | | *Raphanus raphanistrum* | 0.28 ± 0.05 | **0.0004** | 0.39 ± 0.07 | **0.0001** | 0.14 ± 0.09 | 0.1675 |
| | | *Rapistrum rugosum* | 0.28 ± 0.05 | **0.0004** | 0.39 ± 0.07 | **0.0001** | 0.14 ± 0.09 | 0.1675 |
| | | *Sinapis arvensis* | 0.45 ± 0.14 | **< 0.0001** | 0.48 ± 0.25 | **< 0.0001** | 0.60 ± 0.13 | **< 0.0001** |

**Table S6.** The number of path coefficients obtained when fitting the optimised path model on each of the different subsets, which are not contained within the 95% confidence interval around the estimation obtained when fitting the model on the full dataset. The total number of causal relations in the model is 25.

| **Year** | **Plant species** | **Number of paths** |
|---|---|---|
| 2017 & 2018 | *Brassica nigra* | 11 |
| | *Raphanus raphanistrum* | 14 |
| | *Rapistrum rugosum* | 14 |
| | *Sinapis arvensis* | 16 |
| 2017 | All | 11 |
| | *Brassica nigra* | 5 |
| | *Raphanus raphanistrum* | 7 |
| | *Rapistrum rugosum* | 7 |
| | *Sinapis arvensis* | 6 |
| 2018 | All | 9 |
| | *Brassica nigra* | 2 |
| | *Raphanus raphanistrum* | 13 |
| | *Rapistrum rugosum* | 13 |
| | *Sinapis arvensis* | 8 |

**Table S7.** Overview of the direct, indirect, and total effect of variables included in the optimized descriptive piecewise path model on seed production of plants in the open field experiment, calculated for the different subsets of the data based on plant species, year, or plant species by year combination. The volume of plants and the number of reproductive branches were square-root-transformed, and the abundances of folivores, florivores and seed predators were transformed using a log(x+1) transformation.

| | | Early-season number of leaves | | | Early-season volume | | | Early-season herbivore richness | | | Early-season herbivore abundance | | |
|---|---|---|---|---|---|---|---|---|---|---|---|---|---|
| **Year** | **Plant species** | **Direct** | **Indirect** | **Total** | **Direct** | **Indirect** | **Total** | **Direct** | **Indirect** | **Total** | **Direct** | **Indirect** | **Total** |
| 2017 & 2018 | *Brassica nigra* | 0.00 | 0.04 | 0.04 | -0.08 | 0.12 | 0.04 | 0.00 | 0.00 | 0.00 | 0.00 | 0.04 | 0.04 |
| | *Raphanus raphanistrum* | 0.00 | 0.06 | 0.06 | 0.18 | -0.05 | 0.13 | 0.00 | 0.00 | 0.00 | 0.00 | 0.05 | 0.05 |
| | *Sinapis arvensis* | 0.00 | 0.10 | 0.10 | -0.16 | -0.04 | -0.20 | 0.00 | 0.00 | 0.00 | 0.00 | 0.05 | 0.05 |
| | *Rapistrum rugosum* | 0.00 | 0.06 | 0.06 | 0.18 | -0.05 | 0.13 | 0.00 | 0.00 | 0.00 | 0.00 | 0.05 | 0.05 |
| 2017 | All | 0.00 | 0.04 | 0.04 | 0.03 | 0.04 | 0.06 | 0.00 | 0.00 | 0.00 | 0.00 | 0.04 | 0.04 |
| | *Brassica nigra* | 0.00 | 0.09 | 0.09 | -0.09 | 0.17 | 0.08 | 0.00 | 0.00 | 0.00 | 0.00 | 0.04 | 0.04 |
| | *Raphanus raphanistrum* | 0.00 | 0.01 | 0.01 | 0.12 | 0.01 | 0.13 | 0.00 | 0.00 | 0.00 | 0.00 | 0.03 | 0.03 |
| | *Sinapis arvensis* | 0.00 | 0.09 | 0.09 | -0.01 | -0.08 | -0.09 | 0.00 | 0.00 | 0.00 | 0.00 | 0.07 | 0.07 |
| | *Rapistrum rugosum* | 0.00 | 0.01 | 0.01 | 0.12 | 0.01 | 0.13 | 0.00 | 0.00 | 0.00 | 0.00 | 0.03 | 0.03 |
| 2018 | All | 0.00 | 0.08 | 0.08 | -0.14 | 0.09 | -0.05 | 0.00 | 0.00 | 0.00 | 0.00 | 0.11 | 0.11 |
| | *Brassica nigra* | 0.00 | 0.03 | 0.03 | 0.06 | 0.13 | 0.19 | 0.00 | 0.00 | 0.00 | 0.00 | 0.01 | 0.01 |
| | *Raphanus raphanistrum* | 0.00 | 0.06 | 0.06 | -0.06 | 0.09 | 0.03 | 0.00 | 0.00 | 0.00 | 0.00 | 0.13 | 0.13 |
| | *Sinapis arvensis* | 0.00 | 0.18 | 0.18 | -0.08 | -0.03 | -0.11 | 0.00 | 0.00 | 0.00 | 0.00 | -0.05 | -0.05 |
| | *Rapistrum rugosum* | 0.00 | 0.06 | 0.06 | -0.06 | 0.09 | 0.03 | 0.00 | 0.00 | 0.00 | 0.00 | 0.10 | 0.10 |

**Table S7. Continued**

| | | Mid-season number of leaves | | | Mid-season volume | | | Mid-season herbivore richness | | | Mid-season herbivore abundance | | |
|---|---|---|---|---|---|---|---|---|---|---|---|---|---|
| **Year** | **Plant species** | **Direct** | **Indirect** | **Total** | **Direct** | **Indirect** | **Total** | **Direct** | **Indirect** | **Total** | **Direct** | **Indirect** | **Total** |
| 2017 & 2018 | *Brassica nigra* | 0.17 | 0.00 | 0.17 | 0.32 | 0.00 | 0.32 | 0.00 | 0.00 | 0.00 | 0.00 | 0.00 | 0.00 |
| | *Raphanus raphanistrum* | 0.10 | 0.02 | 0.12 | 0.21 | 0.13 | 0.34 | 0.00 | -0.0425 | -0.04 | 0.00 | 0.00 | 0.00 |
| | *Sinapis arvensis* | 0.09 | 0.20 | 0.29 | 0.27 | 0.19 | 0.46 | 0.00 | -0.02 | -0.02 | 0.00 | 0.00 | 0.00 |
| | *Rapistrum rugosum* | 0.10 | 0.02 | 0.12 | 0.21 | 0.13 | 0.34 | 0.00 | -0.04 | -0.04 | 0.00 | 0.00 | 0.00 |
| 2017 | All | 0.10 | 0.02 | 0.12 | 0.38 | 0.04 | 0.42 | 0.00 | -0.01 | -0.01 | 0.00 | 0.00 | 0.00 |
| | *Brassica nigra* | 0.22 | 0.01 | 0.23 | 0.31 | 0.01 | 0.32 | 0.00 | 0.00 | 0.00 | 0.00 | 0.00 | 0.00 |
| | *Raphanus raphanistrum* | -0.02 | 0.05 | 0.03 | 0.14 | 0.18 | 0.32 | 0.00 | -0.07 | -0.07 | 0.00 | 0.00 | 0.00 |
| | *Sinapis arvensis* | 0.10 | 0.20 | 0.30 | 0.21 | 0.20 | 0.41 | 0.00 | 0.03 | 0.03 | 0.00 | 0.00 | 0.00 |
| | *Rapistrum rugosum* | -0.02 | 0.05 | 0.03 | 0.14 | 0.18 | 0.32 | 0.00 | -0.07 | -0.07 | 0.00 | 0.00 | 0.00 |
| 2018 | All | 0.33 | -0.01 | 0.32 | 0.36 | -0.03 | 0.33 | 0.00 | 0.00 | 0.00 | 0.00 | 0.00 | 0.00 |
| | *Brassica nigra* | 0.17 | 0.01 | 0.18 | 0.20 | -0.04 | 0.16 | 0.00 | 0.01 | 0.01 | 0.00 | 0.00 | 0.00 |
| | *Raphanus raphanistrum* | 0.09 | 0.04 | 0.13 | 0.32 | 0.01 | 0.33 | 0.00 | -0.02 | -0.02 | 0.00 | 0.00 | 0.00 |
| | *Sinapis arvensis* | 0.24 | 0.32 | 0.56 | -0.01 | 0.10 | 0.09 | 0.00 | -0.08 | -0.08 | 0.00 | 0.00 | 0.00 |
| | *Rapistrum rugosum* | 0.09 | 0.04 | 0.13 | 0.32 | 0.01 | 0.33 | 0.00 | -0.02 | -0.02 | 0.00 | 0.00 | 0.00 |

**Table S7. Continued**

| Year | Plant species | Reproductive branches | | | Florivore & seed predator abundance | | | Florivore & seed predator richness | | |
|---|---|---|---|---|---|---|---|---|---|---|
| | | Direct | Indirect | Total | Direct | Indirect | Total | Direct | Indirect | Total |
| 2017 & 2018 | *Brassica nigra* | 0.05 | 0.00 | 0.05 | 0.00 | 0.00 | 0.00 | 0.00 | 0.00 | 0.00 |
| | *Raphanus raphanistrum* | 0.01 | 0.00 | 0.01 | 0.00 | 0.00 | 0.00 | 0.00 | 0.00 | 0.00 |
| | *Sinapis arvensis* | 0.28 | 0.00 | 0.28 | 0.00 | 0.00 | 0.00 | 0.00 | 0.00 | 0.00 |
| | *Rapistrum rugosum* | 0.45 | 0.00 | 0.45 | 0.00 | 0.00 | 0.00 | 0.00 | 0.00 | 0.00 |
| 2017 | All | 0.28 | 0.00 | 0.28 | 0.00 | 0.00 | 0.00 | 0.00 | 0.00 | 0.00 |
| | *Brassica nigra* | 0.13 | 0.00 | 0.13 | 0.00 | 0.00 | 0.00 | 0.00 | 0.00 | 0.00 |
| | *Raphanus raphanistrum* | 0.08 | 0.00 | 0.08 | 0.00 | 0.00 | 0.00 | 0.00 | 0.00 | 0.00 |
| | *Sinapis arvensis* | 0.39 | 0.00 | 0.39 | 0.00 | 0.00 | 0.00 | 0.00 | 0.00 | 0.00 |
| | *Rapistrum rugosum* | 0.48 | 0.00 | 0.48 | 0.00 | 0.00 | 0.00 | 0.00 | 0.00 | 0.00 |
| 2018 | All | 0.39 | 0.00 | 0.39 | 0.00 | 0.00 | 0.00 | 0.00 | 0.00 | 0.00 |
| | *Brassica nigra* | -0.09 | 0.00 | -0.09 | 0.00 | 0.00 | 0.00 | 0.00 | 0.00 | 0.00 |
| | *Raphanus raphanistrum* | -0.13 | 0.00 | -0.13 | 0.00 | 0.00 | 0.00 | 0.00 | 0.00 | 0.00 |
| | *Sinapis arvensis* | 0.14 | 0.00 | 0.14 | 0.00 | 0.00 | 0.00 | 0.00 | 0.00 | 0.00 |
| | *Rapistrum rugosum* | 0.60 | 0.00 | 0.60 | 0.00 | 0.00 | 0.00 | 0.00 | 0.00 | 0.00 |

**Table S8.** Overview of effect size of early-season herbivory (Treatment), year of the field season (Year), and their interaction (Treatment*Year) on the different variables related to plant development and the associated herbivore community as used in the path model. Models were adjusted to account for heterogeneity of model residuals across different treatments, across different years, for all factor levels, or were not adjusted when no heterogeneity was found. Values in bold indicate significant factor effects ($p < 0.05$). The volume of plants and the number of reproductive branches were square-root-transformed, and the abundances of folivores, florivores and seed predators were transformed using a log(x+1) transformation.

| Plant species | Parameter | Variance functions | Explanatory variable | $\chi^2$ | df | p-value |
|---|---|---|---|---|---|---|
| *Brassica nigra* | Early season volume | Year | Treatment | 1.2165 | 2 | 0.5443 |
| | | | Year | 154.9821 | 1 | **< 0.0001** |
| | | | Treatment*Year | 1.3904 | 2 | 0.4990 |
| | Early season number of leaves | Year | Treatment | 1.2431 | 2 | 0.5371 |
| | | | Year | 86.9023 | 1 | **< 0.0001** |
| | | | Treatment*Year | 0.5309 | 2 | 0.7669 |
| | Early season abundance | Year | Treatment | 2.3208 | 2 | 0.3134 |
| | | | Year | 0.6540 | 1 | 0.4187 |
| | | | Treatment*Year | 1.2378 | 2 | 0.5385 |
| | Early season richness | | Treatment | 2.1591 | 2 | 0.3398 |
| | | | Year | 5.5717 | 1 | **0.0183** |
| | | | Treatment*Year | 1.1105 | 2 | 0.5739 |
| | Mid-season volume | Year | Treatment | 2.0216 | 2 | 0.3639 |
| | | | Year | 25.8394 | 1 | **< 0.0001** |
| | | | Treatment*Year | 0.3622 | 2 | 0.8344 |
| | Mid-season number of leaves | Year | Treatment | 0.1077 | 2 | 0.9476 |
| | | | Year | 55.8356 | 1 | **< 0.0001** |
| | | | Treatment*Year | 5.7467 | 2 | 0.0565 |
| | Mid-season richness | Year | Treatment | 1.3060 | 2 | 0.5205 |
| | | | Year | 3.0561 | 1 | 0.0804 |
| | | | Treatment*Year | 0.2990 | 2 | 0.8611 |
| | Mid-season abundance | NA | Treatment | 2.5793 | 2 | 0.2754 |
| | | | Year | 2.0115 | 1 | 0.1561 |
| | | | Treatment*Year | 0.1496 | 2 | 0.9279 |
| | Reproductive branches | Treatment | Treatment | 0.9627 | 2 | 0.6180 |
| | | | Year | 25.0070 | 1 | **< 0.0001** |
| | | | Treatment*Year | 0.8323 | 2 | 0.6596 |
| | Florivore and seed predator abundance | NA | Treatment | 0.9471 | 2 | 0.6228 |
| | | | Year | 38.4027 | 1 | **0.0000** |
| | | | Treatment*Year | 2.9212 | 2 | 0.2321 |
| | Florivore and seed predator richness | Year*Treatment | Treatment | 0.6364 | 2 | 0.7275 |
| | | | Year | 0.2451 | 1 | 0.6206 |
| | | | Treatment*Year | 3.2416 | 2 | 0.1977 |
| *Raphanus raphanistrum* | Early season volume | Treatment*Year | Treatment | 0.4248 | 2 | 0.8086 |
| | | | Year | 275.4069 | 1 | **< 0.0001** |
| | | | Treatment*Year | 0.2587 | 2 | 0.8787 |
| | Early season number of leaves | Year | Treatment | 0.2436 | 2 | 0.8853 |
| | | | Year | 245.4692 | 1 | **< 0.0001** |
| | | | Treatment*Year | 0.0779 | 2 | 0.9618 |
| | Early season abundance | NA | Treatment | 5.4497 | 2 | 0.0656 |
| | | | Year | 0.0041 | 1 | 0.9490 |
| | | | Treatment*Year | 0.9890 | 2 | 0.6099 |

**Table S8. Continued**

| Plant species | Parameter | Variance functions | Explanatory variable | $\chi^2$ | df | p-value |
|---|---|---|---|---|---|---|
| *Raphanus raphanistrum* | Early season richness | Treatment*Year | Treatment | 9.4785 | 2 | **0.0087** |
| | | | Year | 4.2343 | 1 | **0.0396** |
| | | | Treatment*Year | 0.0180 | 2 | 0.9910 |
| | Mid-season volume | Year | Treatment | 5.4516 | 2 | 0.0655 |
| | | | Year | 52.3512 | 1 | **< 0.0001** |
| | | | Treatment*Year | 0.3291 | 2 | 0.8483 |
| | Mid-season number of leaves | Year | Treatment | 1.5734 | 2 | 0.4554 |
| | | | Year | 29.1383 | 1 | **< 0.0001** |
| | | | Treatment*Year | 0.1818 | 2 | 0.9131 |
| | Mid-season richness | Treatment*Year | Treatment | 0.0186 | 2 | 0.9907 |
| | | | Year | 11.8175 | 1 | **0.0006** |
| | | | Treatment*Year | 0.0805 | 2 | 0.9606 |
| | Mid-season abundance | Treatment | Treatment | 0.3051 | 2 | 0.8585 |
| | | | Year | 3.6336 | 1 | 0.0566 |
| | | | Treatment*Year | 0.0415 | 2 | 0.9795 |
| | Reproductive branches | Treatment | Treatment | 0.2309 | 2 | 0.8910 |
| | | | Year | 43.0726 | 1 | **< 0.0001** |
| | | | Treatment*Year | 0.2218 | 2 | 0.8950 |
| | Florivore and seed predator abundance | NA | Treatment | 1.1141 | 2 | 0.5729 |
| | | | Year | 91.6850 | 1 | **< 0.0001** |
| | | | Treatment*Year | 1.4600 | 2 | 0.4819 |
| | Florivore and seed predator richness | Year | Treatment | 0.2898 | 2 | 0.8651 |
| | | | Year | 13.7833 | 1 | **0.0002** |
| | | | Treatment*Year | 1.0629 | 2 | 0.5877 |
| *Sinapis arvensis* | Early season volume | Treatment*Year | Treatment | 1.6196 | 2 | 0.4450 |
| | | | Year | 12.8629 | 1 | **0.0003** |
| | | | Treatment*Year | 0.3808 | 2 | 0.8266 |
| | Early season number of leaves | Year | Treatment | 4.5818 | 2 | 0.1012 |
| | | | Year | 13.7504 | 1 | **0.0002** |
| | | | Treatment*Year | 0.9615 | 2 | 0.6183 |
| | Early season abundance | Year | Treatment | 2.0866 | 2 | 0.3523 |
| | | | Year | 0.5903 | 1 | 0.4423 |
| | | | Treatment*Year | 1.6034 | 2 | 0.4486 |
| | Early season richness | Treatment*Year | Treatment | 0.7579 | 2 | 0.6846 |
| | | | Year | 0.0171 | 1 | 0.8959 |
| | | | Treatment*Year | 1.4059 | 2 | 0.4951 |
| | Mid-season volume | Year | Treatment | 7.0779 | 2 | **0.0290** |
| | | | Year | 77.4952 | 1 | **< 0.0001** |
| | | | Treatment*Year | 2.3185 | 2 | 0.3137 |
| | Mid-season number of leaves | Year | Treatment | 0.1803 | 2 | 0.9138 |
| | | | Year | 0.5259 | 1 | 0.4684 |
| | | | Treatment*Year | 2.4153 | 2 | 0.2989 |
| | Mid-season richness | Treatment*Year | Treatment | 1.1451 | 2 | 0.5641 |
| | | | Year | 0.7701 | 1 | 0.3802 |
| | | | Treatment*Year | 1.2341 | 2 | 0.5395 |
| | Mid-season abundance | Year | Treatment | 1.5352 | 2 | 0.4641 |
| | | | Year | 0.8009 | 1 | 0.3708 |
| | | | Treatment*Year | 0.9682 | 2 | 0.6163 |
| | Reproductive branches | Treatment | Treatment | 1.4744 | 2 | 0.4784 |
| | | | Year | 12.2462 | 1 | **0.0005** |
| | | | Treatment*Year | 7.0812 | 2 | **0.0290** |

**Table S8. Continued**

| Plant species | Parameter | Variance functions | Explanatory variable | $\chi^2$ | df | p-value |
|---|---|---|---|---|---|---|
| *Sinapis arvensis* | Florivore and seed predator abundance | Year | Treatment | 2.8423 | 2 | 0.2414 |
| | | | Year | 118.0519 | 1 | **< 0.0001** |
| | | | Treatment*Year | 0.2324 | 2 | 0.8903 |
| | Florivore and seed predator richness | Treatment*Year | Treatment | 1.5139 | 2 | 0.4691 |
| | | | Year | 17.7477 | 1 | **< 0.0001** |
| | | | Treatment*Year | 0.7825 | 2 | 0.6762 |
| *Rapistrum rugosum* | Early season volume | NA | Treatment | 1.0504 | 2 | 0.5914 |
| | | | Year | 21.6738 | 1 | **< 0.0001** |
| | | | Treatment*Year | 0.0355 | 2 | 0.9824 |
| | Early season number of leaves | Treatment*Year | Treatment | 0.0274 | 2 | 0.9864 |
| | | | Year | 60.6256 | 1 | **< 0.0001** |
| | | | Treatment*Year | 0.2705 | 2 | 0.8735 |
| | Early season abundance | NA | Treatment | 2.5489 | 2 | 0.2796 |
| | | | Year | 3.7088 | 1 | 0.0541 |
| | | | Treatment*Year | 0.7698 | 2 | 0.6805 |
| | Early season richness | NA | Treatment | 2.5352 | 2 | 0.2815 |
| | | | Year | 2.5714 | 1 | 0.1088 |
| | | | Treatment*Year | 0.4147 | 2 | 0.8127 |
| | Mid-season volume | Year | Treatment | 3.2889 | 2 | 0.1931 |
| | | | Year | 12.9156 | 1 | **0.0003** |
| | | | Treatment*Year | 5.2741 | 2 | 0.0716 |
| | Mid-season number of leaves | Treatment*Year | Treatment | 2.6034 | 2 | 0.2721 |
| | | | Year | 33.5237 | 1 | **< 0.0001** |
| | | | Treatment*Year | 2.0805 | 2 | 0.3534 |
| | Mid-season richness | Treatment | Treatment | 0.6275 | 2 | 0.7307 |
| | | | Year | 8.8490 | 1 | **0.0029** |
| | | | Treatment*Year | 0.7583 | 2 | 0.6845 |
| | Mid-season abundance | NA | Treatment | 2.3445 | 2 | 0.3097 |
| | | | Year | 4.8593 | 1 | **0.0275** |
| | | | Treatment*Year | 0.7179 | 2 | 0.6984 |
| | Reproductive branches | Year | Treatment | 0.4646 | 2 | 0.7927 |
| | | | Year | 15.7713 | 1 | **< 0.0001** |
| | | | Treatment*Year | 1.2520 | 2 | 0.5347 |
| | Florivore and seed predator abundance | Treatment | Treatment | 2.5345 | 2 | 0.2816 |
| | | | Year | 33.6340 | 1 | **< 0.0001** |
| | | | Treatment*Year | 4.2205 | 2 | 0.1212 |
| | Florivore and seed predator richness | NA | Treatment | 2.4599 | 2 | 0.2923 |
| | | | Year | 5.9864 | 1 | **0.0144** |
| | | | Treatment*Year | 1.5861 | 2 | 0.4525 |

**Table S9.** Overview of the effect size as estimated by generalized additive mixed models of early-season herbivory (Treatment), year of the field season (Year), and their interaction (Treatment*Year) on herbivore species richness, herbivore abundance, the volume of plants, and the number of true leaves observed for the different plant species. Species richness was modelled using a negative binomial probability distribution with log link function, herbivore abundance was modelled using a gaussian probability distribution with an exponential variance function correcting for heterogeneity of residuals over time since the start of the experiment, and the volume of plants and the number of true leaves were modelled using a gamma probability distribution with log link function. The non-linear relationship between the response variables and day since the start of the experiment was estimated by a general smoothing function (General), or by a model estimating a different relationship for each of the two years (Year) or each of the three treatments (Treatment). Models which assumed a different relationship for each year by treatment combination were never selected as most parsimonious based on AIC. In addition to the analysis on the data including both field seasons, we refitted models on each of the two field seasons separately. Values in bold indicate significant factor effects ($p < 0.05$).

| Response | Plant species | Year | Smoothing function | Explanatory variable | $\chi^2$ | df | p-value |
|---|---|---|---|---|---|---|---|
| Richness | *Brassica nigra* | 2017 & 2018 | Year | Treatment | 1.6813 | 2 | 0.4314 |
| | | | | Year | 4.7127 | 1 | **0.0299** |
| | | | | Treatment * Year | 0.0225 | 2 | 0.9888 |
| | | 2017 | General | Treatment | 0.3580 | 2 | 0.8361 |
| | | 2018 | General | Treatment | 1.2422 | 2 | 0.5373 |
| | *Raphanus raphanistrum* | 2017 & 2018 | Year | Treatment | 1.6093 | 2 | 0.4472 |
| | | | | Year | 0.5455 | 1 | 0.4602 |
| | | | | Treatment * Year | 0.0038 | 2 | 0.9981 |
| | | 2017 | General | Treatment | 0.7103 | 2 | 0.7011 |
| | | 2018 | General | Treatment | 0.8388 | 2 | 0.6574 |
| | *Sinapis arvensis* | 2017 & 2018 | Year | Treatment | 0.5315 | 2 | 0.7666 |
| | | | | Year | 8.1448 | 1 | **0.0043** |
| | | | | Treatment * Year | 3.5903 | 2 | 0.1661 |
| | | 2017 | General | Treatment | 1.8020 | 2 | 0.4062 |
| | | 2018 | General | Treatment | 2.7441 | 2 | 0.2536 |
| | *Rapistrum rugosum* | 2017 & 2018 | Year | Treatment | 1.0846 | 2 | 0.5814 |
| | | | | Year | 1.3580 | 1 | 0.2439 |
| | | | | Treatment * Year | 1.8229 | 2 | 0.4019 |
| | | 2017 | General | Treatment | 1.4931 | 2 | 0.4740 |
| | | 2018 | General | Treatment | 1.3007 | 2 | 0.5219 |
| Abundance (log x+1) | *Brassica nigra* | 2017 & 2018 | Year | Treatment | 0.1926 | 2 | 0.9082 |
| | | | | Year | 5.3453 | 1 | **0.0208** |
| | | | | Treatment * Year | 4.2329 | 2 | 0.1205 |
| | | 2017 | General | Treatment | 1.5037 | 2 | 0.4715 |
| | | 2018 | General | Treatment | 4.0201 | 2 | 0.1340 |
| | *Raphanus raphanistrum* | 2017 & 2018 | Year | Treatment | 0.1924 | 2 | 0.9083 |
| | | | | Year | 5.3446 | 1 | **0.0208** |
| | | | | Treatment * Year | 4.2309 | 2 | 0.1206 |
| | | 2017 | General | Treatment | 0.1803 | 2 | 0.9138 |
| | | 2018 | General | Treatment | 1.0278 | 2 | 0.5982 |
| | *Sinapis arvensis* | 2017 & 2018 | Year | Treatment | 1.5002 | 2 | 0.4723 |
| | | | | Year | 17.9710 | 1 | **<0.0001** |
| | | | | Treatment * Year | 9.7089 | 2 | **0.0078** |
| | | 2017 | General | Treatment | 0.9440 | 2 | 0.6237 |
| | | 2018 | General | Treatment | 15.2120 | 2 | **0.0005** |

**Table S9. Continued**

| Response | Plant species | Year | Smoothing functions | Explanatory variable | $\chi^2$ | df | p-value |
|---|---|---|---|---|---|---|---|
| Abundance (log x+1) | *Rapistrum rugosum* | 2017 & 2018 | Year | Treatment | 0.7506 | 2 | 0.6871 |
| | | | | Year | 0.0424 | 1 | 0.8368 |
| | | | | Treatment * Year | 0.2829 | 2 | 0.8681 |
| | | 2017 | General | Treatment | 1.8728 | 2 | 0.3920 |
| | | 2018 | General | Treatment | 6.2678 | 2 | **0.0436** |
| $\sqrt{\text{Volume}}$ | *Brassica nigra* | 2017 & 2018 | Year | Treatment | 0.2262 | 2 | 0.8931 |
| | | | | Year | 15.267 | 1 | **0.0009** |
| | | | | Treatment * Year | 3.5118 | 2 | 0.1727 |
| | | 2017 | General | Treatment | 0.8518 | 2 | 0.6532 |
| | | 2018 | General | Treatment | 3.4707 | 2 | 0.1763 |
| | *Raphanus raphanistrum* | 2017 & 2018 | Year | Treatment | 1.6965 | 2 | 0.4282 |
| | | | | Year | 17.525 | 1 | **0.0003** |
| | | | | Treatment * Year | 0.7108 | 2 | 0.7009 |
| | | 2017 | General | Treatment | 0.8573 | 2 | 0.6514 |
| | | 2018 | General | Treatment | 1.4009 | 2 | 0.4964 |
| | *Sinapis arvensis* | 2017 & 2018 | Year | Treatment | 3.4006 | 2 | 0.1826 |
| | | | | Year | 1.3582 | 1 | 0.2439 |
| | | | | Treatment * Year | 4.7238 | 2 | 0.9746 |
| | | 2017 | General | Treatment | 0.4422 | 2 | 0.8016 |
| | | 2018 | General | Treatment | 4.385 | 2 | 0.1116 |
| | *Rapistrum rugosum* | 2017 & 2018 | Year | Treatment | 0.8151 | 2 | 0.6653 |
| | | | | Year | 0.2607 | 1 | 0.6096 |
| | | | | Treatment * Year | 7.8847 | 2 | **0.0194** |
| | | 2017 | General | Treatment | 4.2505 | 2 | 0.1194 |
| | | 2018 | General | Treatment | 4.6318 | 2 | 0.0987 |
| Number of leaves | *Brassica nigra* | 2017 & 2018 | Treatment | Treatment | 3.1494 | 2 | 0.2071 |
| | | | | Year | 54.5222 | 1 | **<0.0001** |
| | | | | Treatment * Year | 0.0885 | 2 | 0.9567 |
| | | 2017 | General | Treatment | 10.177 | 2 | **0.0062** |
| | | 2018 | General | Treatment | 3.6609 | 2 | 0.1603 |
| | *Raphanus raphanistrum* | 2017 & 2018 | Year | Treatment | 0.0262 | 2 | 0.9870 |
| | | | | Year | 6.1129 | 1 | **0.0134** |
| | | | | Treatment * Year | 1.8193 | 2 | 0.4027 |
| | | 2017 | General | Treatment | 0.6101 | 2 | 0.7371 |
| | | 2018 | General | Treatment | 1.1109 | 2 | 0.5738 |
| | *Sinapis arvensis* | 2017 & 2018 | Year | Treatment | 0.3311 | 2 | 0.8474 |
| | | | | Year | 0.6265 | 1 | 0.4286 |
| | | | | Treatment * Year | 1.9504 | 2 | 0.3771 |
| | | 2017 | General | Treatment | 0.0740 | 2 | 0.9637 |
| | | 2018 | General | Treatment | 0.3678 | 2 | 0.8320 |
| | *Rapistrum rugosum* | 2017 & 2018 | Year | Treatment | 0.1188 | 2 | 0.9895 |
| | | | | Year | 0.2855 | 1 | 0.5931 |
| | | | | Treatment * Year | 2.4221 | 2 | 0.2979 |
| | | 2017 | General | Treatment | 0.7213 | 2 | 0.6972 |
| | | 2018 | General | Treatment | 2.4575 | 2 | 0.2927 |

**Figure S1.** Observed herbivore abundance since start of the experiment (expressed in days) on *Sinapis arvensis* plants in 2017 (panel A) and 2018 (panel B). Plants were challenged by *Myzus persicae* aphids (green), *Pieris rapae* caterpillars (blue), or were left untreated (orange). Individual observations are represented by dots. The solid line represents the average abundance of herbivores as estimated by a generalized additive model, along with the standard error (shaded area) and 95% confidence interval (outlined by the dashed lines) around this estimate. The GAMM estimated a different relation between the abundance of herbivores and day since the start of the experiment for each of the two years.

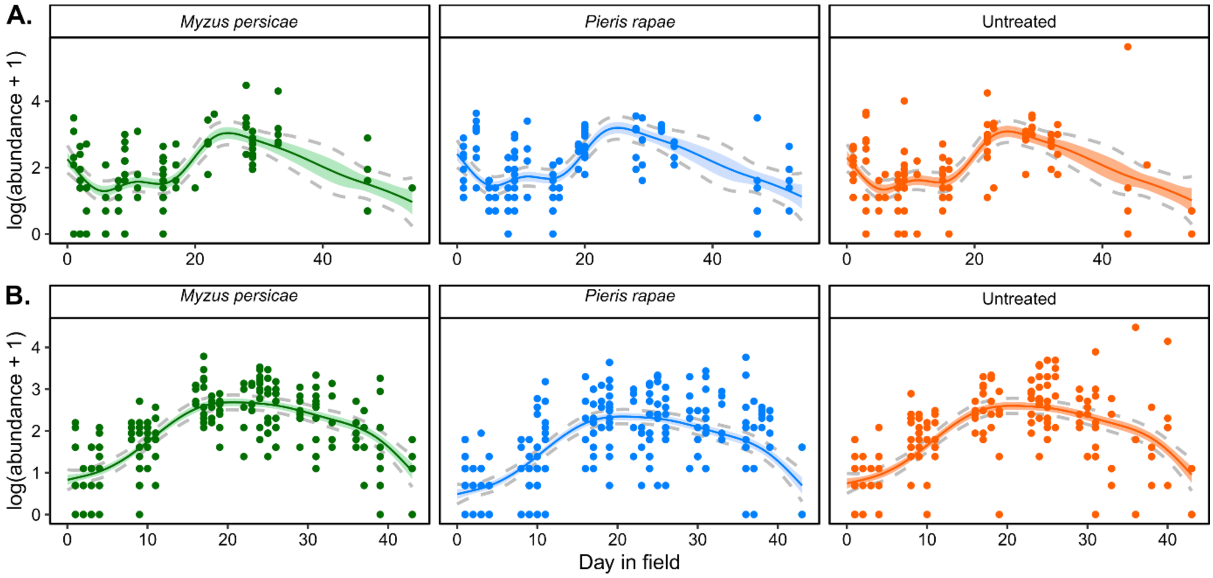


**Figure S2.** Observed volume of *Rapistrum rugosum* plants since start of the experiment (expressed in days) in 2017 (panel A) and 2018 (panel B). Plants were challenged by *Myzus persicae* aphids (green), *Pieris rapae* caterpillars (blue), or were left untreated (orange). Individual observations are represented by dots. The solid line represents the average abundance of herbivores as estimated by a generalized additive model, along with the standard error (shaded area) and 95% confidence interval (outlined by the dashed lines) around this estimate. The GAMM estimated a different relation between the volume of plants and day since the start of the experiment for each of the two years.

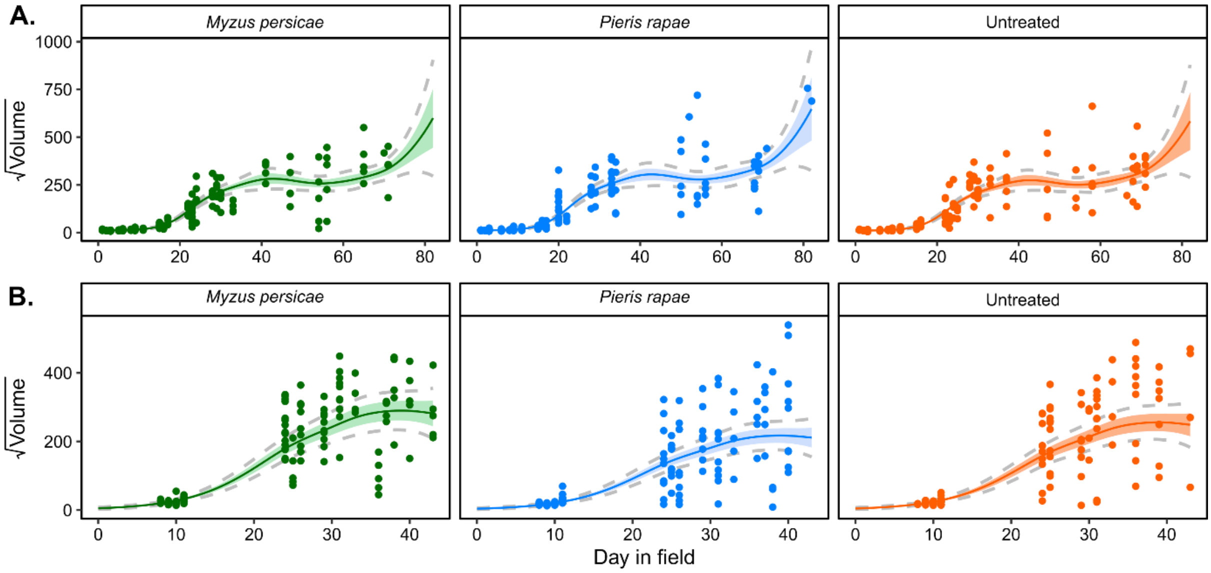

**Figure S3.** Observed herbivore abundance since start of the experiment (expressed in days) on *Rapistrum rugosum* plants in 2017 (panel A) and 2018 (panel B). Plants were challenged by *Myzus persicae* aphids (green), *Pieris rapae* caterpillars (blue), or were left untreated (orange). Individual observations are represented by dots. The solid line represents the average abundance of herbivores as estimated by a generalized additive model, along with the standard error (shaded area) and 95% confidence interval (outlined by the dashed lines) around this estimate. The GAMM estimated a different relation between the abundance of herbivores and day since the start of the experiment for each of the two years.

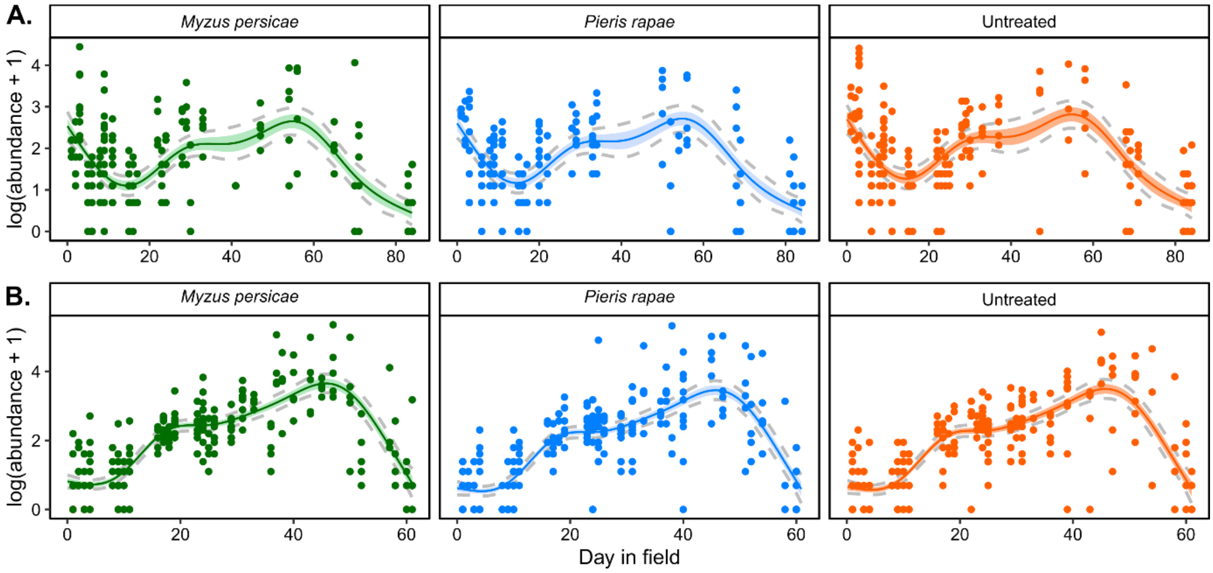


**Figure S4.** Observed number of true leaves of *Brassica nigra* plants since start of the experiment (expressed in days) in 2017 (panel A) and 2018 (panel B). Plants were challenged by *Myzus persicae* aphids (green), *Pieris rapae* caterpillars (blue), or were left untreated (orange). Individual observations are represented by dots. The solid line represents the average abundance of herbivores as estimated by a generalized additive model, along with the standard error (shaded area) and 95% confidence interval (outlined by the dashed lines) around this estimate. The GAMM estimated a different relation between the number of true leaves and day since the start of the experiment for each of the two years.

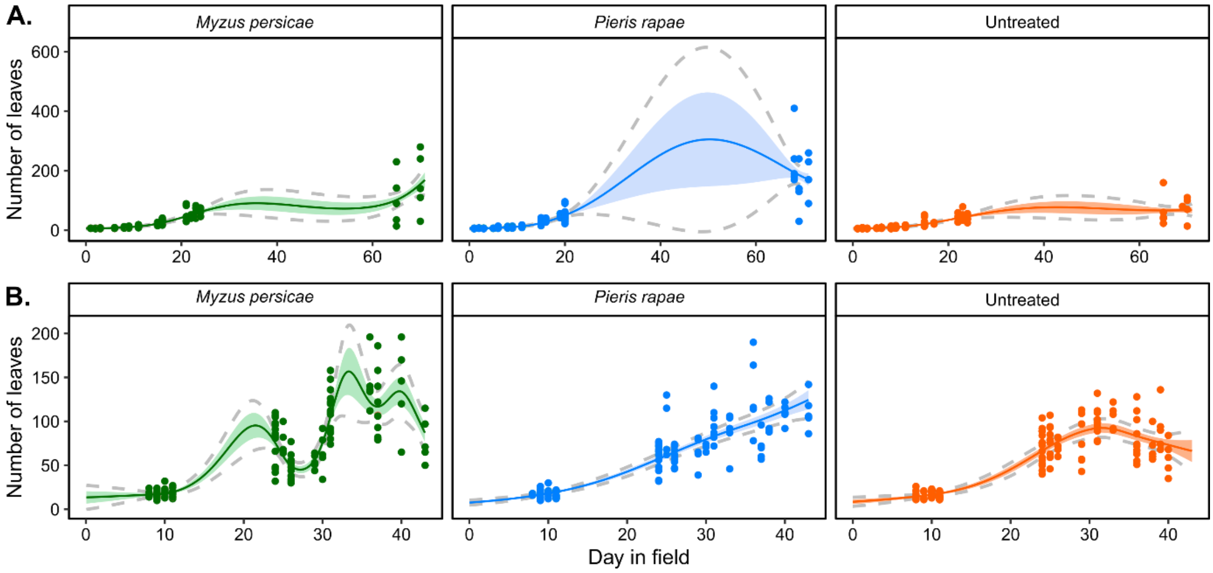

**Figure S5.** Ordination of observed herbivore community composition (expressed by incidence of herbivores, panels A and B), and structure (expressed by Hellinger-transformed herbivore abundance data, panels C and D) in *Brassica nigra* according to three NMDS ordination axes (stress = 0.18 and 0.15 respectively). Plants were challenged by *Myzus persicae* aphids (green), *Pieris rapae* caterpillars (blue), or were left untreated (orange). Coloured shapes represent the centroid of the variation in communities associated with plants belonging to the different treatment – year combinations. Error bars around the centroids represent the 95% distribution of plants of each treatment in the multivariate space. Ellipses are coloured according to the year of the field season and depict the 95% interval of a multivariate t-distribution around the centroids of all plants in each of the field season, independent of their treatment.

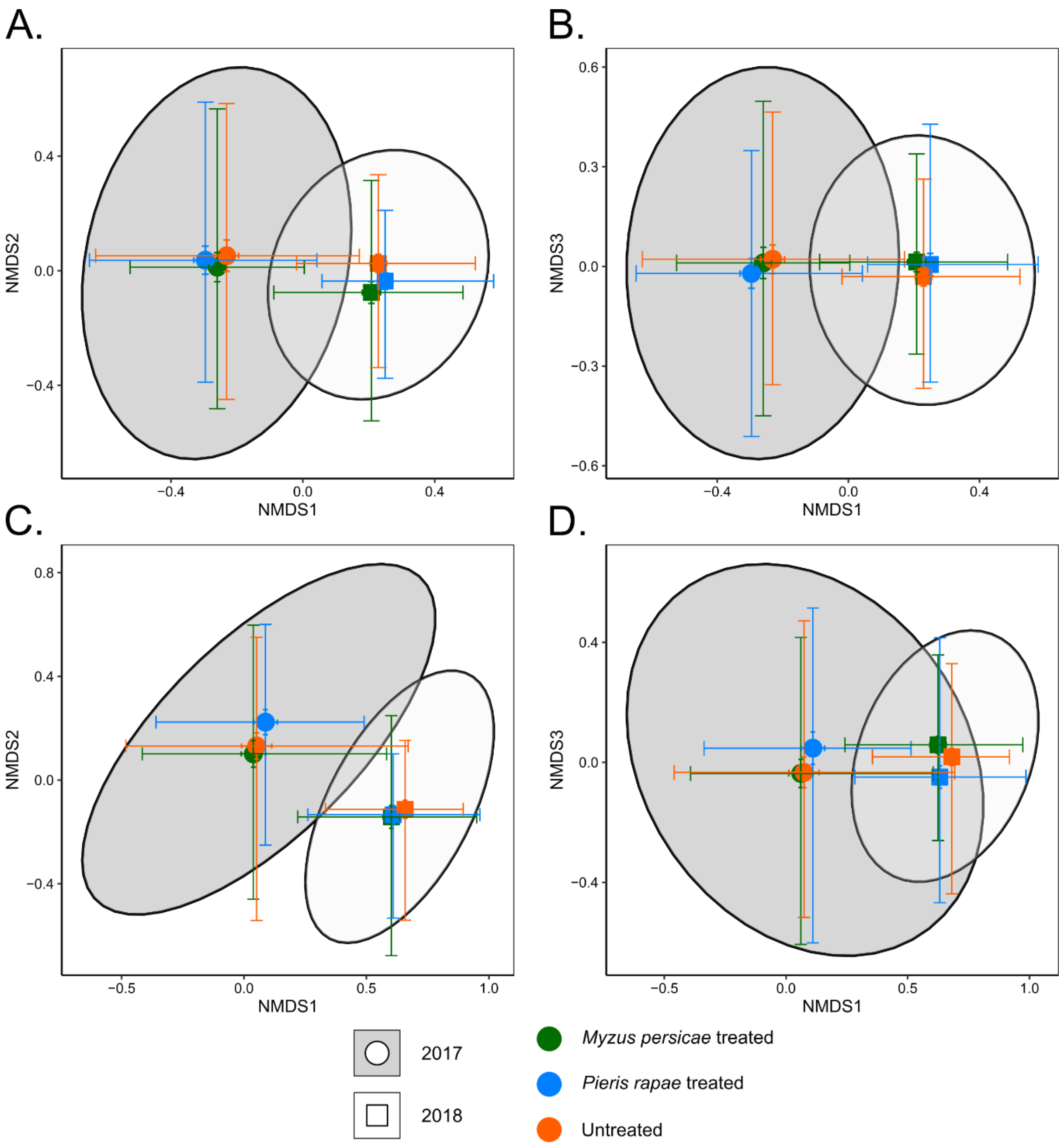

**Figure S6.** Ordination of observed herbivore community composition (expressed by incidence of herbivores, panels A and B), and structure (expressed by Hellinger-transformed herbivore abundance data, panels C and D) in *Raphanus raphanistrum* according to three NMDS ordination axes (stress = 0.18 and 0.16 respectively). Plants were challenged by *Myzus persicae* aphids (green), *Pieris rapae* caterpillars (blue), or were left untreated (orange). Coloured shapes represent the centroid of the variation in communities associated with plants belonging to the different treatment – year combinations. Error bars around the centroids represent the 95% distribution of plants of each treatment in the multivariate space. Ellipses are coloured according to the year of the field season and depict the 95% interval of a multivariate t-distribution around the centroids of all plants in each of the field season, independent of their treatment.

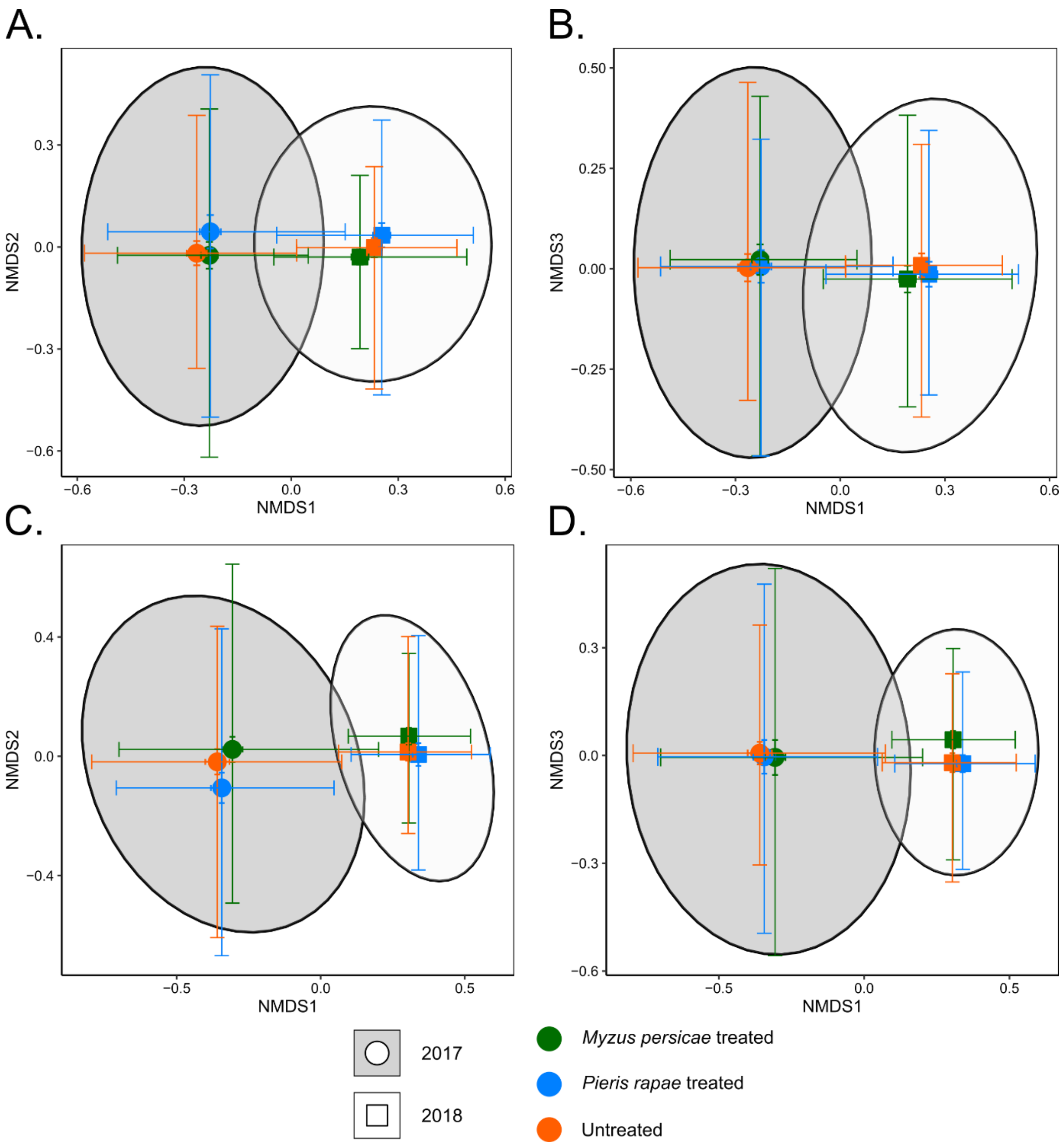

**Figure S7.** Ordination of observed herbivore community composition (expressed by incidence of herbivores, panels A and B), and structure (expressed by Hellinger-transformed herbivore abundance data, panels C and D) in *Sinapis arvensis* according to three NMDS ordination axes (stress = 0.17 and 0.17 respectively). Plants were challenged by *Myzus persicae* aphids (green), *Pieris rapae* caterpillars (blue), or were left untreated (orange). Coloured shapes represent the centroid of the variation in communities associated with plants belonging to the different treatment – year combinations. Error bars around the centroids represent the 95% distribution of plants of each treatment in the multivariate space. Ellipses are coloured according to the year of the field season and depict the 95% interval of a multivariate t-distribution around the centroids of all plants in each of the field season, independent of their treatment.

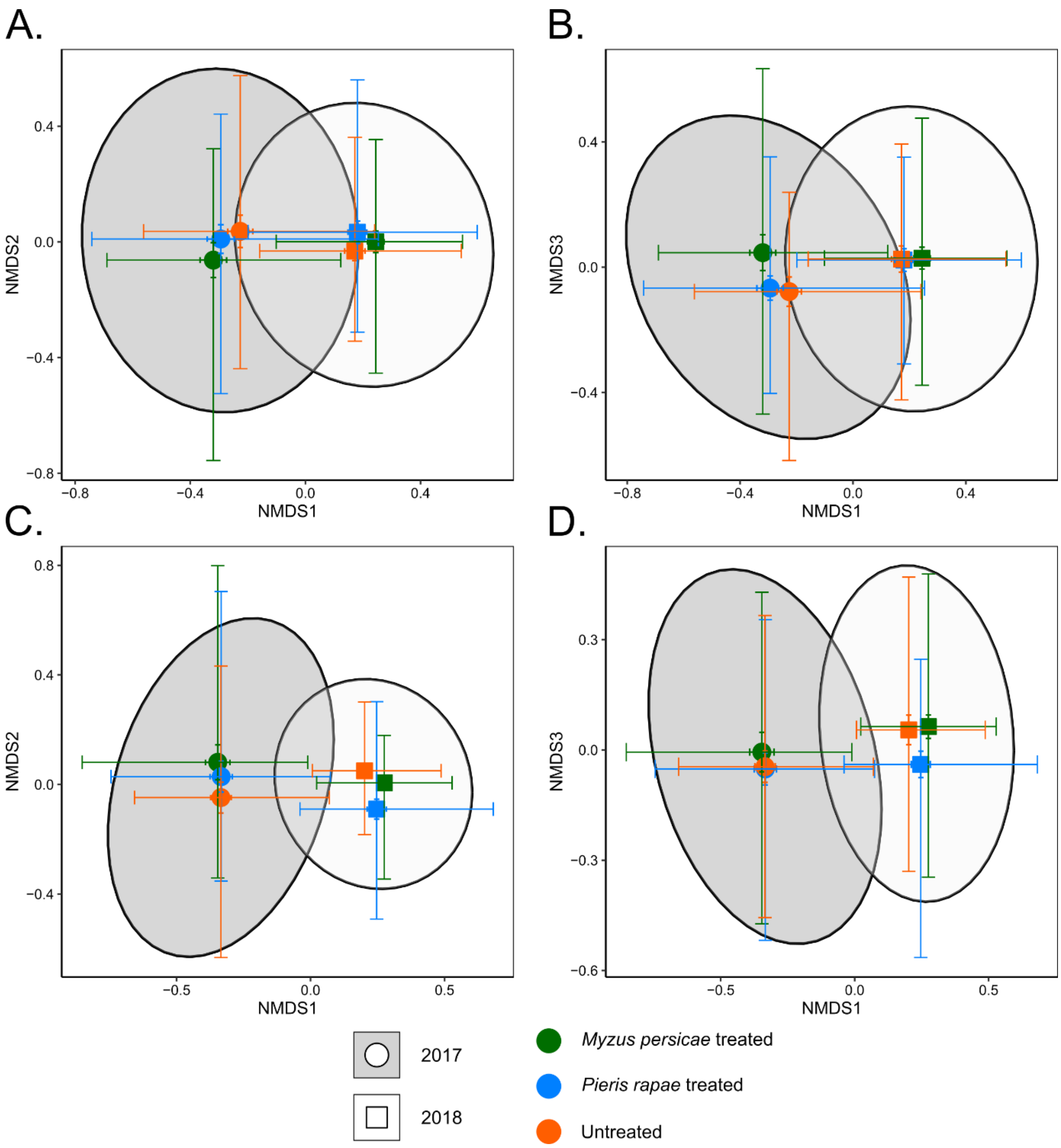

**Figure S8.** Ordination of observed herbivore community composition (expressed by incidence of herbivores, panels A and B), and structure (expressed by Hellinger-transformed herbivore abundance data, panels C and D) in *Rapistrum rugosum* according to three NMDS ordination axes (stress = 0.18 and 0.16 respectively). Plants were challenged by *Myzus persicae* aphids (green), *Pieris rapae* caterpillars (blue), or were left untreated (orange). Coloured shapes represent the centroid of the variation in communities associated with plants belonging to the different treatment – year combinations. Error bars around the centroids represent the 95% distribution of plants of each treatment in the multivariate space. Ellipses are coloured according to the year of the field season and depict the 95% interval of a multivariate t-distribution around the centroids of all plants in each of the field season, independent of their treatment.

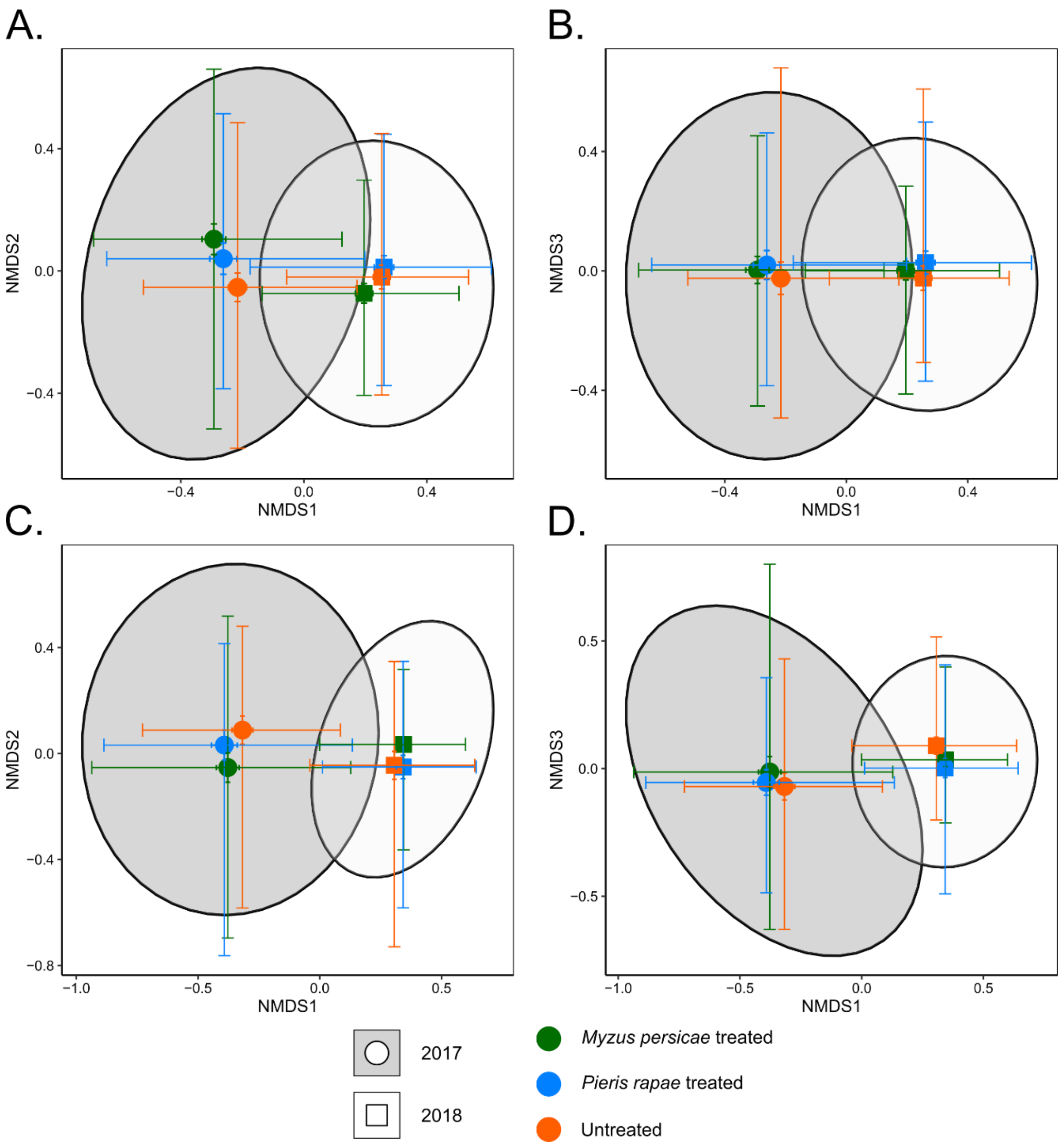

**Table S10.** Results of the PERMANOVA analysis testing the effects of early-season herbivory treatments on the composition (incidence) and structure (weighted abundance, calculated as Hellinger-transformed herbivore abundance data) of the full herbivore community associated with individual plants in each plant species by year combination. To account for dependency of observations, we applied a stratified permutation design (1000 permutations) with 100 rounds of random sampling to ensure equal replication across treatment levels. The table presents the quantile distribution of pseudo-F values with associated $R^2$ and significance. Values in bold indicate significant treatment effects ($p < 0.05$)

| | | | Composition | | | | Structure | | | |
|---|---|---|---|---|---|---|---|---|---|---|
| **Year** | **Plant species** | **Percentile** | **Pseudo - F** | **$R^2$** | **df** | **p-value** | **Pseudo - F** | **$R^2$** | **df** | **p-value** |
| 2017 | *Brassica nigra* | 5 | 0.5582 | 0.0167 | 2 | 0.966 | 0.5644 | 0.0168 | 2 | 0.988 |
| | | 25 | 0.8404 | 0.0248 | 2 | 0.883 | 0.7774 | 0.0230 | 2 | 0.945 |
| | | 50 | 1.0689 | 0.0314 | 2 | 0.776 | 0.9502 | 0.0280 | 2 | 0.877 |
| | | 75 | 1.2725 | 0.0371 | 2 | 0.645 | 1.0862 | 0.0319 | 2 | 0.752 |
| | | 95 | 1.4636 | 0.0424 | 2 | 0.630 | 1.3447 | 0.0392 | 2 | 0.624 |
| | *Raphanus raphanistrum* | 5 | 0.6145 | 0.0176 | 2 | 0.935 | 0.7393 | 0.0210 | 2 | 0.967 |
| | | 25 | 0.8714 | 0.0247 | 2 | 0.798 | 0.8787 | 0.0248 | 2 | 0.913 |
| | | 50 | 1.0915 | 0.0306 | 2 | 0.698 | 1.0112 | 0.0285 | 2 | 0.868 |
| | | 75 | 1.4608 | 0.0405 | 2 | 0.461 | 1.2300 | 0.0344 | 2 | 0.739 |
| | | 95 | 1.8754 | 0.0514 | 2 | 0.279 | 1.4390 | 0.0400 | 2 | 0.498 |
| | *Sinapis arvensis* | 5 | 0.9037 | 0.0342 | 2 | 0.727 | 0.8911 | 0.0338 | 2 | 0.810 |
| | | 25 | 1.2077 | 0.0453 | 2 | 0.592 | 1.1506 | 0.0432 | 2 | 0.807 |
| | | 50 | 1.5090 | 0.0561 | 2 | 0.297 | 1.3173 | 0.0491 | 2 | 0.586 |
| | | 75 | 1.8564 | 0.0678 | 2 | 0.142 | 1.5336 | 0.0567 | 2 | 0.439 |
| | | 95 | 2.3464 | 0.0842 | 2 | 0.101 | 1.7281 | 0.0635 | 2 | 0.357 |
| | *Rapistrum rugosum* | 5 | 0.7453 | 0.0221 | 2 | 0.888 | 0.6681 | 0.0198 | 2 | 0.956 |
| | | 25 | 1.1337 | 0.0333 | 2 | 0.563 | 0.8074 | 0.0239 | 2 | 0.926 |
| | | 50 | 1.3271 | 0.0386 | 2 | 0.554 | 0.9601 | 0.0283 | 2 | 0.917 |
| | | 75 | 1.6060 | 0.0462 | 2 | 0.377 | 1.1482 | 0.0336 | 2 | 0.785 |
| | | 95 | 2.2303 | 0.0633 | 2 | 0.129 | 1.3574 | 0.0395 | 2 | 0.677 |
| 2018 | *Brassica nigra* | 5 | 0.6017 | 0.0171 | 2 | 0.838 | 0.6797 | 0.0193 | 2 | 0.886 |
| | | 25 | 0.8954 | 0.0253 | 2 | 0.726 | 0.8725 | 0.0247 | 2 | 0.609 |
| | | 50 | 1.1522 | 0.0323 | 2 | 0.592 | 0.9765 | 0.0275 | 2 | 0.600 |
| | | 75 | 1.3681 | 0.0381 | 2 | 0.396 | 1.1013 | 0.0309 | 2 | 0.472 |
| | | 95 | 1.6310 | 0.0451 | 2 | 0.194 | 1.4435 | 0.0402 | 2 | 0.295 |
| | *Raphanus raphanistrum* | 5 | 0.6155 | 0.0175 | 2 | 0.911 | 1.1676 | 0.0327 | 2 | 0.635 |
| | | 25 | 1.0418 | 0.0293 | 2 | 0.751 | 1.3869 | 0.0386 | 2 | 0.495 |
| | | 50 | 1.2424 | 0.0348 | 2 | 0.578 | 1.5722 | 0.0436 | 2 | 0.406 |
| | | 75 | 1.5864 | 0.0440 | 2 | 0.349 | 1.8700 | 0.0514 | 2 | 0.133 |
| | | 95 | 2.0033 | 0.0549 | 2 | 0.138 | 2.4842 | 0.0672 | 2 | 0.082 |
| | *Sinapis arvensis* | 5 | 0.4985 | 0.0142 | 2 | 0.936 | 1.3926 | 0.0388 | 2 | 0.382 |
| | | 25 | 0.7610 | 0.0216 | 2 | 0.823 | 1.7340 | 0.0479 | 2 | 0.207 |
| | | 50 | 1.0001 | 0.0282 | 2 | 0.635 | 2.0557 | 0.0562 | 2 | 0.058 |
| | | 75 | 1.2083 | 0.0338 | 2 | 0.529 | 2.3850 | 0.0647 | 2 | **0.026** |
| | | 95 | 1.5399 | 0.0427 | 2 | 0.404 | 2.8400 | 0.0761 | 2 | **0.008** |
| | *Rapistrum rugosum* | 5 | 0.3889 | 0.0111 | 2 | 0.997 | 0.5074 | 0.0145 | 2 | 0.992 |
| | | 25 | 0.6306 | 0.0180 | 2 | 0.934 | 0.7425 | 0.0211 | 2 | 0.909 |
| | | 50 | 0.9129 | 0.0258 | 2 | 0.881 | 0.9629 | 0.0272 | 2 | 0.775 |
| | | 75 | 1.0589 | 0.0298 | 2 | 0.687 | 1.1838 | 0.0332 | 2 | 0.471 |
| | | 95 | 1.3932 | 0.0388 | 2 | 0.574 | 1.4310 | 0.0398 | 2 | 0.353 |
